# Attenuated, Not Absent. Trusting Autonomous Systems Tracks Reliability More Closely, With Less Neural Engagement, Than Trusting Humans

**DOI:** 10.64898/2026.11.14.766654

**Authors:** Daniel A. Rogers, Alexander Weuthen, Anthony Finn, Matthias Schlesewsky, Ina Bornkessel-Schlesewsky, Markus Ullsperger

## Abstract

Users of autonomous systems and AI assistants decide whether to accept a system’s output, yet whether such active trust decisions engage the same evaluative architecture as trust between people remains unknown. Forty-four participants made trial-by-trial decisions to trust or distrust a human or autonomous system partner during fMRI, with identical prescripted accuracy for both partners across five difficulty levels. Despite equivalent performance, trust in humans exceeded their reliability. At the highest difficulty level, the estimated probability of trusting the human partner was 72% but only 51% for the system, despite both performing at 52% accuracy. Distrust decisions recruited a prespecified performance monitoring network of valuation, salience, social-perceptual and affective regions. On hard trials, this network remained functionally coupled for both partners. Across the task, trust and distrust were discriminable in every component for both, with greater distrust-related activation and discriminability for human partners. Discriminability differences were expressed on hard trials, where partner accuracy was uncertain and trust diverged between partners. These findings indicate that trust in humans and autonomous systems engages a shared evaluative architecture, but that engagement is attenuated during autonomous system interaction. Whereas our earlier passive monitoring study found the P3a to system errors absent with no behavioural difference, an active decision appears to impose a floor of engagement, so that technology-related cognitive economization, a reduction in cognitive and social-affective resources for non-human partners, takes the form of attenuation rather than absence. Alongside this attenuation, trust in the system tracked its reliability without exceeding it.

---

When Lee and See (2004) laid the foundations for the study of trust in automation, the neurological level was identified as crucial for a complete account, yet two decades later it remains the least developed. Trust in automation is a problem of reliance under uncertainty, wherein operators of autonomous systems in fields from medical diagnostics to industrial supervision constantly monitor system performance and decide to accept or override system outputs (Hancock et al., 2021), a decision that users of AI assistants built on large language models now make routinely. These are value-based decisions that weigh partner performance against expected outcomes and gains against social risks (Rilling & Sanfey, 2011). Such decisions recruit a domain-general performance monitoring network that evaluates outcomes, detects expectation violations, and initiates adaptive behaviour (Ullsperger et al., 2014a) and is thought to integrate signals from cognitive and affective brain regions (Ullsperger et al., 2014b). Trust decisions between people recruit elements of this same broad evaluative network (Bellucci et al., 2017), while trust has also been suggested as dependent on interactions among large-scale neural networks (Krueger & Meyer-Lindenberg, 2019). Though neuroimaging has investigated trust in technological partners, it is unknown whether trust in automated systems recruits a similar network of brain regions as trust in humans.

Prior fMRI work has compared human and machine agents at the level of individual regions, or of data-driven connectivity among task-activated areas (Aimone et al., 2014; Goodyear et al., 2016, 2017; Hopko et al., 2024; Krach et al., 2008), but has not tested whether an autonomous partner engages the evaluative network specified by performance monitoring theory. Characterising trust in autonomous systems as a network therefore requires first establishing the components of that network in the case where they are best understood, trust between people. Within the performance monitoring framework, the network comprises functionally distinct components that each contribute specific properties to a unified evaluative response. Valuation regions, including the orbitofrontal cortex and ventral striatum, encode subjective value and outcome predictions (O’Doherty, 2004; Rolls et al., 2020); in trust paradigms, multiround decisions consistently recruit the ventral striatum, consistent with reward prediction and learning a partner’s reputation across repeated exchange (Bellucci et al., 2017). Salience regions, including the anterior insula and anterior midcingulate cortex, signal uncertainty, conflict, and the need for behavioural revision (Craig, 2009; Uddin, 2015). In trust and related exchange games, anterior insula activation is consistently observed, particularly in one-shot decisions, and is linked to aversive affect, expectancy of norm compliance, and sensitivity to surprising outcomes (Aimone et al., 2014; Bellucci et al., 2017; Bellucci et al., 2018). Social-perceptual processing is supported by the right posterior superior temporal sulcus, which responds more strongly to interactive than non-interactive behaviour, encodes whether interactions are cooperative or competitive, and supports attribution of intentionality (Isik et al., 2017; Lahnakoski et al., 2012; Walbrin et al., 2018). The amygdala contributes an affective and social-risk component. Selective basolateral amygdala damage abolishes adaptive updating of investments in a repeated trust game (Rosenberger et al., 2019), and primate basolateral neurons encode social preferences and reward to self and others during social choice (Chang et al., 2015; see Gangopadhyay et al., 2021, for a review). Integrated within the performance monitoring network, these components together evaluate the trustworthiness of human partners.

While the regions supporting trust decisions in humans are well characterised, emerging evidence suggests this architecture may respond differently depending on partner type. Neural responses to artificial agents differ from responses to human partners, with modulation of social motivation and mentalising regions reported across several paradigms (Pfeiffer et al., 2014; Wiese et al., 2017). Network-level comparison of human-robot and human-human interaction has found differences at perceptual and language levels without a comparable difference in the theory-of-mind network, indicating that artificial and human partners can produce distinct network profiles (Hogenhuis & Hortensius, 2022). Further work shows amygdala responses are greater for human compared to technological partners (Rauchbauer et al., 2019), and can predict rejection of artificial agents (Rosenthal-von der Pütten et al., 2019). In prior work, errors by an autonomous system partner elicited no P3a, an index of frontal attentional engagement, although performance evaluation signals to those errors were preserved, whereas errors by a human partner elicited a P3a (Rogers et al., 2026). We interpreted this as technology-related cognitive economization (TRCE), a reduction in the cognitive and social-affective resources allocated to a non-human partner. Behaviour did not differ between partners in that study. Whether partner-related differences persist under active decision conditions remains unclear. When individuals must actively evaluate and decide whether to trust or distrust a partner’s performance, task demands may impose a baseline level of engagement that alters how such effects are expressed.

An active decision also allows trust to be compared with the partner’s actual reliability. Trust in automation is appropriate when it is calibrated to the system’s reliability (Lee & See, 2004), and miscalibration has been reported in both directions, as reliance inflated by expectations of machine perfection (Dzindolet et al., 2003) and as algorithms abandoned more readily than humans after the same error (Dietvorst et al., 2015). Whether a difference in trust between human and automated partners reflects miscalibrated trust in the machine or in the person is most directly tested when both partners perform identically and trust is compared with that performance.

In addition, the role of uncertainty may be influential. As partner difficulty increases and partner outcomes become less predictable, the contribution of affective and social information to decision-making may change. Difficulty and cognitive effort modulate salience and performance monitoring regions that are also implicated in trust. Perceptual decision difficulty increases anterior insula activation (Lamichhane et al., 2016), and cognitive effort alters connectivity between dorsal anterior cingulate cortex and task-relevant cortical areas (Aben et al., 2020). When the uncertainty concerns another agent, anterior cingulate cortex tracks how far that agent’s information should guide choice (Behrens et al., 2008). Whether difficulty alters how partner type influences these systems during active trust decisions remains unexplored.

The present study addressed these questions using functional magnetic resonance imaging (fMRI) during a trust decision task involving either a human or autonomous system partner across varying levels of difficulty. We examined activation using multi-level general linear models (GLM) and, within predefined functional networks, representational structure using multivariate pattern analysis (MVPA) and coupling using beta-series functional connectivity to test whether these regions operate as a coupled network during trust decisions. Based on prior work, we expected that partner type would modulate neural processing within the performance monitoring network, and that partner difficulty would influence both behavioural and neural responses. Because both partners performed with identical accuracy, we could also test whether trust in each partner was calibrated to its actual reliability. By combining activation, representational, and connectivity approaches, this study aimed to characterise whether neural processing differs between human and autonomous system interaction, and how and under what conditions such differences emerge.

## Methods

### Participants

Forty-four adults aged 20 to 38 years (28 male and 16 female; 1 left-handed) participated in the study. All had normal or corrected-to-normal vision, reported no history of neurological or psychiatric disorder and no MRI contraindications. Participants were pre-screened by telephone and provided written informed consent prior to testing. The study was approved by the ethics committee of Otto-von-Guericke University. Participants received €10 per hour for participating and a €1.50 performance bonus was awarded for any task runs in which a participant earned at least 8,000 points (rounded up to €5.00 if bonus was achieved in all three runs). Participants were allocated between-subjects to either a human partner (*n* = 22) or autonomous system partner (*n* = 22) condition. Two participants were excluded from the multivariate pattern and beta-series connectivity analyses. Neither exclusion resulted from motion thresholds or any other criterion applied to otherwise valid data. Rather, they were due to data-integrity failures identified during single-trial estimation. Sub-01 was excluded because stimulus blocks used a non-standard run structure, meaning the data were not compatible with single-trial models of the remaining participants. Sub-35 was excluded because the single-trial model failed to converge. Both participants remained valid for inclusion in GLM and behaviour analyses (see Supplementary Materials for full rationale of participant exclusion). Multivariate and connectivity analyses therefore included 42 participants (human *n* = 20, system *n* = 22).

### Experimental Design and Task

The task used a trust decision paradigm in which participants made image classifications while tracking the performance of a partner and making decisions to trust or distrust the partner’s classification of the same image. The task was programmed in Presentation (Neurobehavioral Systems) and presented in German. Participants completed three runs, each of which included five progressive difficulty levels with 25 trials per level for a total trial count of 125 trials per run and 375 trials per participant. All images were of cats or dogs taken from the Microsoft Kaggle Dogs versus Cats dataset (Microsoft, 2017) and were overlaid with a transparent diagonal black and white mask. The masking was used to manipulate the difficulty of discriminating the image as a cat or a dog for the participants themselves while also providing a plausible explanation for the varied level of partner performance. Higher masking therefore reduced the evidence available for classification, increasing uncertainty in the participant’s own decision. Unique target images were used for each trial so that participants only saw each image once. The task structure is summarised in Figure 1.

**Figure 1.**
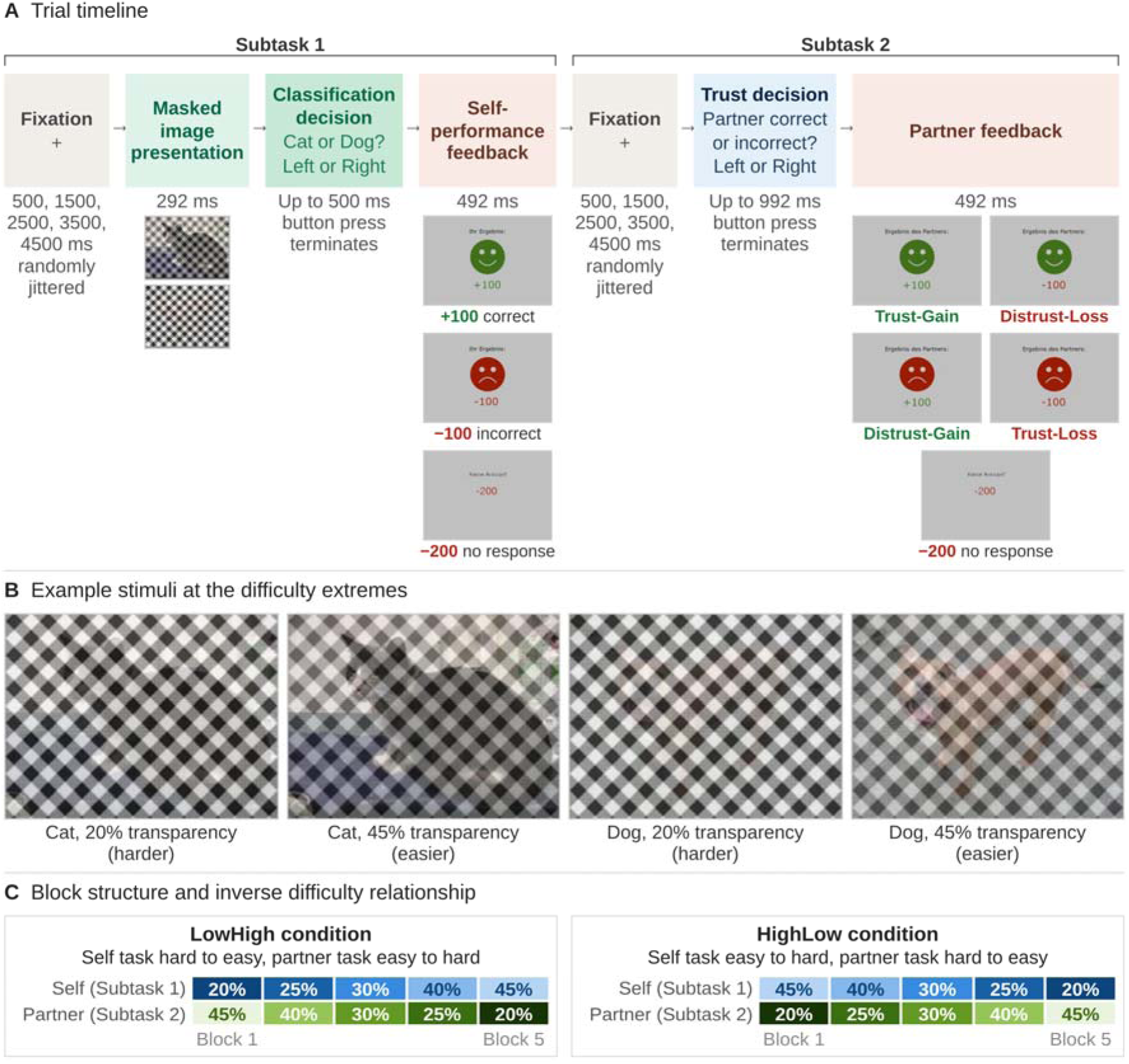
Task Structure for the Nemesys Paradigm. *Note.* (A) Trial timeline showing the sequence and duration of each phase of Subtask 1 and Subtask 2. Each feedback screen appears beneath the phase in which it was shown, with correct, incorrect and missed responses at self-performance feedback, and the four partner feedback outcomes (Trust-Gain, Distrust-Loss, Distrust-Gain and Trust-Loss) and missed trust decisions at partner feedback. (B) Example masked stimuli at the hardest (20% transparency) and easiest (45% transparency) difficulty levels. (C) Block structure for the two difficulty-order conditions, showing the inverse relationship between self-performance difficulty and partner difficulty across the five blocks of each run. German screen text: Ihr Ergebnis = your result; Ergebnis des Partners = partner’s result; Keine Antwort = no response.

#### Subtask 1: Self-Performance Classification

On each trial, participants were presented with a photograph of a cat or a dog that had been obscured by a black-and-white mask. The mask varied in transparency across five difficulty levels (labelled 20, 25, 30, 40, and 45 in the task, corresponding to the percentage transparency of the mask, so that lower values were more heavily masked and harder to discriminate). Participants indicated their classification as quickly as possible via a button press, with the left button for cat, right button for dog. Correct responses earned +100 points; incorrect responses resulted in −100 points; a failure to respond within the time limit incurred a −200-point penalty (displayed as “Keine Antwort!” in German which translates to “No Answer!” with −200 in red text). Outcome feedback for Subtask 1 was displayed as a smiley face (green, upturned mouth) paired with the points earned (+100 in green), or a frowning face (red, downturned mouth) paired with the points lost (−100 in red), under the heading “Ihr Ergebnis:” (Your result). See Figure 1 for a full task and trial overview.

#### Subtask 2: Partner Trust Decision

Immediately following the self-performance feedback on each trial a time-jittered fixation cross was presented, after which participants predicted whether their assigned partner would respond correctly or incorrectly on the same trial. They were told that their partner had previously performed the identical task, but with difficulty that was the inverse of their own. When the participant’s masking level was light (easy), the partner’s masking level was heavy (hard), and vice versa (see Figure 1 for an overview of the difficulty manipulation). This inverse difficulty relationship meant that partner performance could be anticipated to change systematically across difficulty levels, providing a structured basis for participants to be able to learn to trust or distrust their partner’s performance across the task. This task manipulation was deliberately implemented to foster a sense of teamwork between the participant and their partner such that the participant should have been able to rely on their partner to perform well when their own self-performance subtask was difficult and vice versa.

Participants made trust decisions on each trial by predicting partner classification outcomes via a button press. Predicting their partner would be correct was a decision to trust the partner on that trial and predicting their partner would be incorrect was a decision to distrust. Immediately following the trust decision partner feedback was then displayed as a smiley or frowning face (identical iconography to self-feedback) under the heading “Ergebnis des Partners:” (Partner’s result), together with the participant’s points earned or lost (±100). The partner feedback’s emotional expression (smiling or frowning) indicated whether the partner had been correct or incorrect, and this was independent of the participant’s prediction outcome. A smiling face with +100 indicated the partner was correct and the prediction was correct (Trust-Gain), whilst a smiling face with −100 indicated the partner was correct but the prediction was incorrect (Distrust-Loss). Analogously, a frowning face with +100 indicated the partner was incorrect and the prediction was correct (Distrust-Gain), and a frowning face with −100 indicated the partner was incorrect and the prediction was incorrect (Trust-Loss).

#### Partner Condition

In the human partner condition, participants were told their partner was another person who had completed the task in an earlier session. In the autonomous system condition, participants were told their partner was an automated classification system. In reality, partner performances were prescripted and task accuracy was identical across partner conditions, with partner identity being a between-subjects manipulation. The partner was correct on 92%, 84%, 76%, 60% and 52% of trials at partner difficulty levels 1 to 5 respectively. All other task parameters were identical across conditions.

#### Difficulty Counterbalancing

Difficulty order was counterbalanced between participants. Within each partner condition, four sub-versions counterbalanced difficulty order (HighLow vs. LowHigh, named for the self-performance image transparency running from high to low or from low to high) and stimulus sequence (versions A and B). In the HighLow condition, mask opacity increased across the five stages of each run (i.e., Subtask 1 became progressively harder), while in the LowHigh condition, mask opacity decreased (Subtask 1 became progressively easier). The inverse applied to the partner. Within each difficulty order, two stimulus sequence versions (A and B) further counterbalanced item presentation to account for the effects of the unique image stimuli used in the experiment. This yielded four partner-by-order sub-conditions (Human-HighLow, Human-LowHigh, System-HighLow, and System-LowHigh), each presented in versions A and B.

### MRI Data Acquisition

MRI data were acquired on a 3-Tesla Siemens Prisma scanner equipped with a 64-channel head coil. Structural images were obtained using a magnetisation-prepared rapid gradient echo (MPRAGE) sequence (voxel size = 1 × 1 × 1 mm; matrix = 192 × 256 × 256).

Functional blood-oxygen-level-dependent (BOLD) images were acquired using a T2*-weighted single-echo gradient-echo echo-planar imaging (EPI) sequence developed at the Center for Magnetic Resonance Research (CMRR), University of Minnesota (cmrr_mbep2d_bold, release R016; Moeller et al., 2010; Xu et al., 2013). Sixty-four oblique axial slices were acquired in interleaved order, tilted 17.1° from the transverse towards the coronal plane, giving whole-brain coverage with 2 mm isotropic voxels and a 10% interslice gap (TR = 2.0 s, TE = 30 ms, flip angle = 80°, field of view = 212 × 212 mm, acquisition matrix = 106 × 106, multiband acceleration factor = 2, in-plane GRAPPA acceleration factor = 2, phase-encoding direction anterior-to-posterior, total readout time = 28.9 ms). Each participant completed three functional runs of the task.

### fMRI Preprocessing

MRI data were converted to Brain Imaging Data Structure (BIDS) format and preprocessed using fMRIPrep 23.0.2 (Esteban et al., 2019; RRID:SCR_016216), which is based on Nipype 1.8.6. The following preprocessing steps were performed for each functional run. A reference volume and skull-stripped version were generated using fMRIPrep’s internal methodology. Six rigid-body head-motion parameters (three translations and three rotations) were estimated relative to the BOLD reference volume using mcflirt (FSL 6.0.5.1; Jenkinson et al., 2002). BOLD runs were slice-timing corrected to 0.959 s (the midpoint of the 0–1.92 s slice acquisition range) using 3dTshift (AFNI; Cox & Hyde, 1997).

The BOLD reference was co-registered to the T1w reference image using mri_coreg (FreeSurfer) followed by boundary-based registration (flirt, FSL 6.0.5.1; Greve & Fischl, 2009; Jenkinson & Smith, 2001), with six degrees of freedom. No susceptibility distortion correction was applied. Volume-based spatial normalisation to MNI152NLin6Asym standard space was performed through nonlinear registration using antsRegistration (ANTs 2.3.3). Functional images were resampled to 2 mm isotropic resolution in MNI space using Lanczos interpolation via antsApplyTransforms.

Framewise displacement (FD; Power et al., 2014) and DVARS were computed for each run. Frames exceeding FD > 0.5 mm or standardised DVARS > 1.5 were flagged as motion outliers by fMRIPrep. The six rigid-body motion parameters were the only confound regressors entered into the first-level models. No motion outlier regressors or CompCor components (Behzadi et al., 2007) were included.

### Behavioural Analyses

Trust decisions were analysed using a generalised linear mixed model with a binomial link function, fitted by maximum likelihood with Laplace approximation (lme4). Trust decisions (Trust/Distrust) were the binary dependent variable, modelled as a function of partner (human/system) and partner difficulty, with their interaction and random intercepts for participant. Difficulty was modelled as a five-level factor. Type II Wald chi-square tests estimated effects, with estimated marginal means and partner contrasts computed on the probability scale for each difficulty level.

### fMRI Analyses: General Linear Model

#### Event Definition

Trial onset times were aligned to the onset of the first scanner pulse of each run and expressed in seconds with event durations modelled as instantaneous (0.005 s). Twelve event types were modelled in four families. Self-performance feedback comprised SelfFBGain, SelfFBLoss, and SelfFBNoChoice for trials on which no self-performance response was registered. The partner trust decision comprised TrustQ, marking the onset of the trust prompt, together with the decision itself modelled as either Trust or Distrust. Partner feedback comprised four event types crossing partner accuracy with the participant’s points outcome, being OtherCorrectGain, OtherCorrectLoss, OtherErrorGain, and OtherErrorLoss. Two fixation events, Fixation1 and Fixation2, completed the model. Event files were formatted to BIDS specification (.tsv). The full first-level design matrix, including the confound regressors described below, is shown in Figure S1.

#### First-Level GLM

First-level GLMs were estimated in Python using nilearn’s FirstLevelModel (Abraham et al., 2014). For each participant, all three runs were modelled simultaneously within a single GLM. The haemodynamic response function (HRF) was modelled using the Glover canonical HRF (Glover, 1999). A cosine drift model was applied for high-pass filtering (0.01 Hz cutoff, equivalent to a 100 s period). Signal scaling was not applied.

Confound regressors included the six rigid-body motion parameters estimated by fMRIPrep (three translations: trans_x, trans_y, trans_z; three rotations: rot_x, rot_y, rot_z). A whole-brain mask derived from the fMRIPrep anatomical segmentation was applied during model fitting.

#### Second-Level GLM

Second-level analyses were conducted using nilearn’s SecondLevelModel. First-level contrast images were spatially smoothed with a 6 mm FWHM Gaussian kernel prior to second-level modelling. The second-level design matrix included a constant regressor (mean across all *N* = 44 participants) and a partner group regressor (Human = +1, System = −1). Three first-level contrasts were carried forward to second level. These were Distrust > Trust, SelfFBLoss > SelfFBGain, and the equivalent loss versus gain contrast for partner feedback, combining OtherCorrectLoss and OtherErrorLoss against OtherCorrectGain and OtherErrorGain. The group mean was computed for all three, and the Human > System partner group difference was computed for Distrust > Trust only. Statistical maps were thresholded using voxel-wise false discovery rate (FDR) correction at *q* < .05.

### Region of Interest Definition

Spherical regions of interest (ROIs) for the multivariate and connectivity analyses were defined in two steps. For each region, a coordinate range was first compiled from prior literature on valuation, salience, social cognition and threat processing, since published coordinates for these regions span a range and do not converge on a single location. Deriving the search range from several studies instead of a single published coordinate follows Poldrack’s (2007) recommendation for ROI specification. Within each literature-defined range, the ROI centre was then placed at the local activation peak in the second-level Distrust > Trust map, computed on the constant regressor across all 44 participants. Locations are shown in Figure 2, and coordinates, sphere radii and peak activation are given in Table 1.

**Figure 2.**
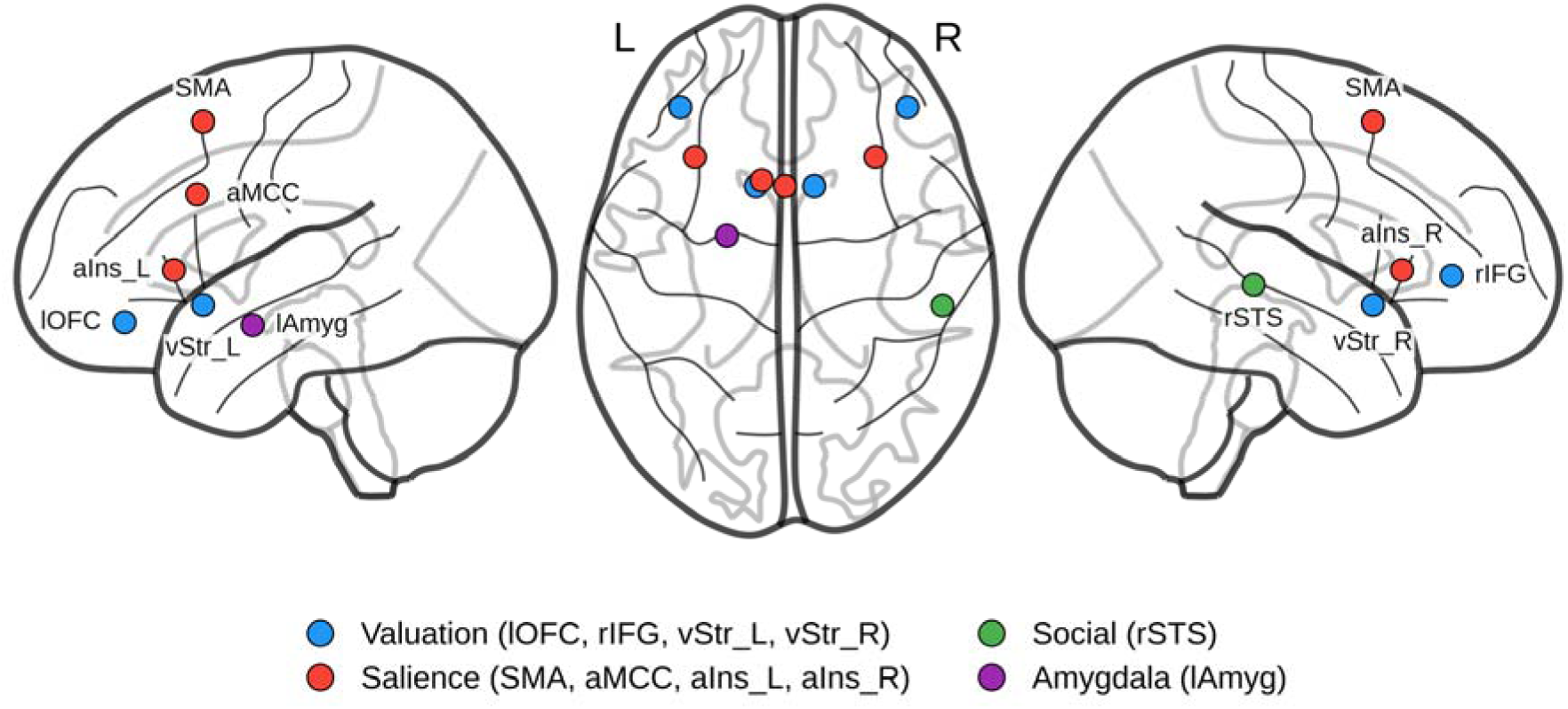
Locations of the Ten Regions of Interest in MNI152 Space, Shown as Glass Brain Projections and Coloured by Network Assignment. *Note.* Regions are labelled in the left and right views. Coordinates and sphere radii are given in Table 1.

**Table 1.** Regions of Interest, Coordinates, Sphere Radii and Peak Activation.

| ROI | Network | MNI (*x*, *y*, *z*) | Radius (mm) | Peak *z* |
|---|---|---|---|---|
| lOFC | Valuation | −36, 40, −12 | 5 | 5.96 |
| rIFG | Valuation | 42, 40, 4 | 5 | 4.72 |
| vStr_L | Valuation | −10, 13, −6 | 4 | 3.99 |
| vStr_R | Valuation | 10, 13, −6 | 4 | 3.83 |
| SMA | Salience | 0, 13, 57 | 6 | 6.80 |
| aMCC | Salience | −8, 15, 32 | 5 | 4.90 |
| aIns_L | Salience | −31, 23, 6 | 5 | 6.58 |
| aIns_R | Salience | 31, 23, 6 | 5 | 6.16 |
| rSTS | Social | 54, −28, 1 | 5 | 5.30 |
| lAmyg | Amygdala | −20, −4, −13 | 4 | 3.41 |

*Note.* Coordinates are in MNI152 space. Each ROI is a sphere of the stated radius centred on the local activation peak within a coordinate range compiled from prior literature. Peak z is the maximum *z* statistic within the sphere in the second-level Distrust > Trust map. lOFC = left lateral orbitofrontal cortex; rIFG = right inferior frontal gyrus; vStr = ventral striatum; SMA = supplementary motor area; aMCC = anterior midcingulate cortex; aIns = anterior insula; rSTS = right superior temporal sulcus; lAmyg = left amygdala; L = left; R = right.

The search ranges were fixed by prior literature and are independent of the present data. ROI centres were placed using the group mean across both partner conditions, which is orthogonal to the human versus system comparison, so the partner effects reported in the decoding and coupling analyses are independent of the selection step (Kriegeskorte et al., 2009).

### Multivariate Pattern Analysis

#### Trial-Level Beta Extraction (Least Squares Separate)

Trial-level activation patterns were estimated using the least squares separate (LSS) approach (Abdulrahman & Henson, 2016; Mumford et al., 2012). Single-trial estimates were computed for the trust and distrust decision trials only. For each decision trial, a separate GLM was estimated containing one regressor for that trial and one nuisance regressor collapsing the remaining decision trials in that run. The HRF was modelled using the Glover canonical HRF (Glover, 1999). A cosine basis set was used for high-pass filtering (0.01 Hz cutoff). Trial events were modelled as instantaneous (duration = 0.005 s), with onset times drawn from the trial event table. Design matrices were constructed using nilearn’s make_first_level_design_matrix, and parameter estimates were obtained via ordinary least squares (scipy.linalg.lstsq; validated against nilearn’s FirstLevelModel, *r* = 1.000).

Confound regressors comprised the six rigid-body motion parameters (trans_x, trans_y, trans_z, rot_x, rot_y, rot_z) extracted from fMRIPrep’s confound files. No spatial smoothing was applied at any stage of the MVPA pipeline. Beta maps for Trust and Distrust decision trials were extracted and saved for each participant and run.

#### Feature Extraction

For each trial, voxel-level beta values within each ROI group were extracted from the corresponding LSS beta map using nilearn’s NiftiSpheresMasker. Features from all ROIs within a group were concatenated into a single multivariate feature vector. Approximate voxel counts per group were: Full network (≈708 voxels), Valuation (228 voxels), Salience (366 voxels), Social/rSTS (81 voxels), and Amygdala (33 voxels). The total trial-level feature dataset comprised approximately 16,019 trials.

#### Classification and Cross-Validation

For each participant, Trust versus Distrust decisions were decoded using a Linear Support Vector Machine (LinearSVC; C = 1.0, class_weight = ‘balanced’; scikit-learn). Classification accuracy was estimated using five-fold stratified cross-validation, reduced to k equal to the minority-class n where fewer than five trials of a class were available, and leave-one-out where fewer than three. Participants were excluded from a specific analysis if all their trials in that subset belonged to a single class (precluding binary classification).

#### Analysis Levels

Two analysis levels were implemented. Level one used all available trials in three subsets (all participants combined, human partner trials only, and system partner trials only). The second level repeated the same all participants, human only and system only comparisons separately by difficulty bins for Easy difficulty (difficulty levels 1 and 2; *n* ≈ 6,408 trials) and Hard difficulty (difficulty levels 4 and 5; *n* ≈ 6,404 trials). Difficulty level 3 (medium) was excluded from the second level analysis to maximise contrast between difficulty extremes. Bins were defined by masking level with behavioural modelling of trust and distrust decisions providing independent support for the approach to binning trials.

Partner differences in trust probability were negligible at the two levels forming the Easy bin (contrasts of .008 and .029, both *p* > .25), smaller at the excluded middle level (.078, *p* = .028), and largest at the two levels forming the Hard bin (.166 and .205, both *p* < .001). As such, partner identity was consequential behaviourally at the levels forming the Hard bin while no measurable consequence was seen in the Easy bin. The behavioural choice data thus provide a basis for the difficulty bin approach that is independent from the neural analysis.

#### Group-Level Inference

For each analysis, participant-level decoding accuracies were tested with one-sample *t* tests against chance (.50; two-tailed; scipy.stats.ttest_1samp). Partner group comparisons used Welch independent-samples *t* tests (scipy.stats.ttest_ind with equal_var = False), as partner condition was manipulated between subjects and variances were unequal. Cohen’s *d* is reported with the pooled standard deviation. Multiple comparisons correction was applied using the Benjamini–Hochberg FDR procedure (Benjamini & Hochberg, 1995), pooling all 60 tests from Levels one and two (significance threshold *q* < .05).

### Beta-Series Functional Connectivity

Beta-series functional connectivity was computed to address two questions (Rissman et al., 2004). The first was whether the ten regions operated as a coupled set or as independent regions. The second was whether a change in that coupling could account for the partner differences observed in the activation and decoding analyses. The same trial-level LSS beta estimates and ten spherical ROIs described above were used in this analysis, meaning no further estimations were required. For each participant, mean betas within ROIs per trial formed a beta series, and connectivity was computed as the Pearson correlation between the beta series of every ROI pair across trials. This provided a 10 × 10 connectivity matrix with 45 unique pairs per participant. All correlations were Fisher *z*-transformed prior to subsequent statistical analysis.

Connectivity was computed within hard trials only, separately for trust and for distrust decisions. Hard trials were selected because this is where the partner effect was located in the decoding analysis and where partner identity was behaviourally consequential. Hard trials were available for every participant in both decision types, with a minimum of 23 trust trials (human median 106, system median 90) and 8 distrust trials (human median 41, system median 60).

Within-group coupling was tested against zero with one-sample *t* tests for each ROI pair, separately for the human group and the system group, in each decision type. Partner group differences used Welch independent-samples *t* tests comparing Fisher *z* values between groups for each pair, within each decision type. The within-participant change from trust to distrust was tested with paired *t* tests in each partner group, and the difference between groups in that change with Welch tests. Every family was corrected across the 45 pairs using the Benjamini–Hochberg false discovery rate procedure (Benjamini & Hochberg, 1995).

### Software

#### Preprocessing and GLM Analyses

Preprocessing was performed using fMRIPrep 23.0.2 (Esteban et al., 2019; RRID:SCR_016216) running on Nipype 1.8.6, which internally used FSL 6.0.5.1, AFNI (Cox & Hyde, 1997), ANTs 2.3.3, FreeSurfer, and Nilearn 0.9.1. First- and second-level GLM analyses were implemented in Python using nilearn (Abraham et al., 2014).

#### MVPA Analyses

MVPA analyses were implemented in Python 3.11 using the following packages: numpy, pandas, nibabel, nilearn (design matrix construction and ROI masking), scikit-learn (LinearSVC classifier, cross-validation), scipy (OLS solver, statistical tests), statsmodels (FDR correction via multipletests), matplotlib and seaborn (visualisation), and openpyxl (Excel export).

## Results

### Trust Is Withdrawn More Readily From the System Partner, in Line With Its Reliability

Trust decisions varied systematically as a function of difficulty and partner type. Partner accuracy was prespecified and matched across partner conditions, so any difference in trust between conditions reflected how participants evaluated the partner, not how the partner performed. The binomial generalised linear mixed model revealed a significant Difficulty × Partner interaction, χ²(4) = 46.57, *p* < .001, indicating that trust declined more steeply across difficulty for system than for human partners. Main effects of difficulty, χ²(4) = 1057.98, *p* < .001, and partner group, χ²(1) = 7.68, *p* = .006, were also significant.

Estimated trust probability declined with difficulty in both conditions and diverged between them as difficulty increased (Figure 3A; Table 2). The partner difference was reliable from level 3 onward. Partner identity therefore had no measurable behavioural consequence while the partner was performing well, and became consequential as partner errors became frequent, with trust withdrawn more readily from the system partner. That withdrawal tracked the partner’s actual accuracy, which fell within the 95% confidence interval of system trust at every difficulty level, whereas trust in the human partner exceeded it from level 2 onward, by 20 percentage points at the highest level (Figure 3B; Table 2).

**Figure 3.**
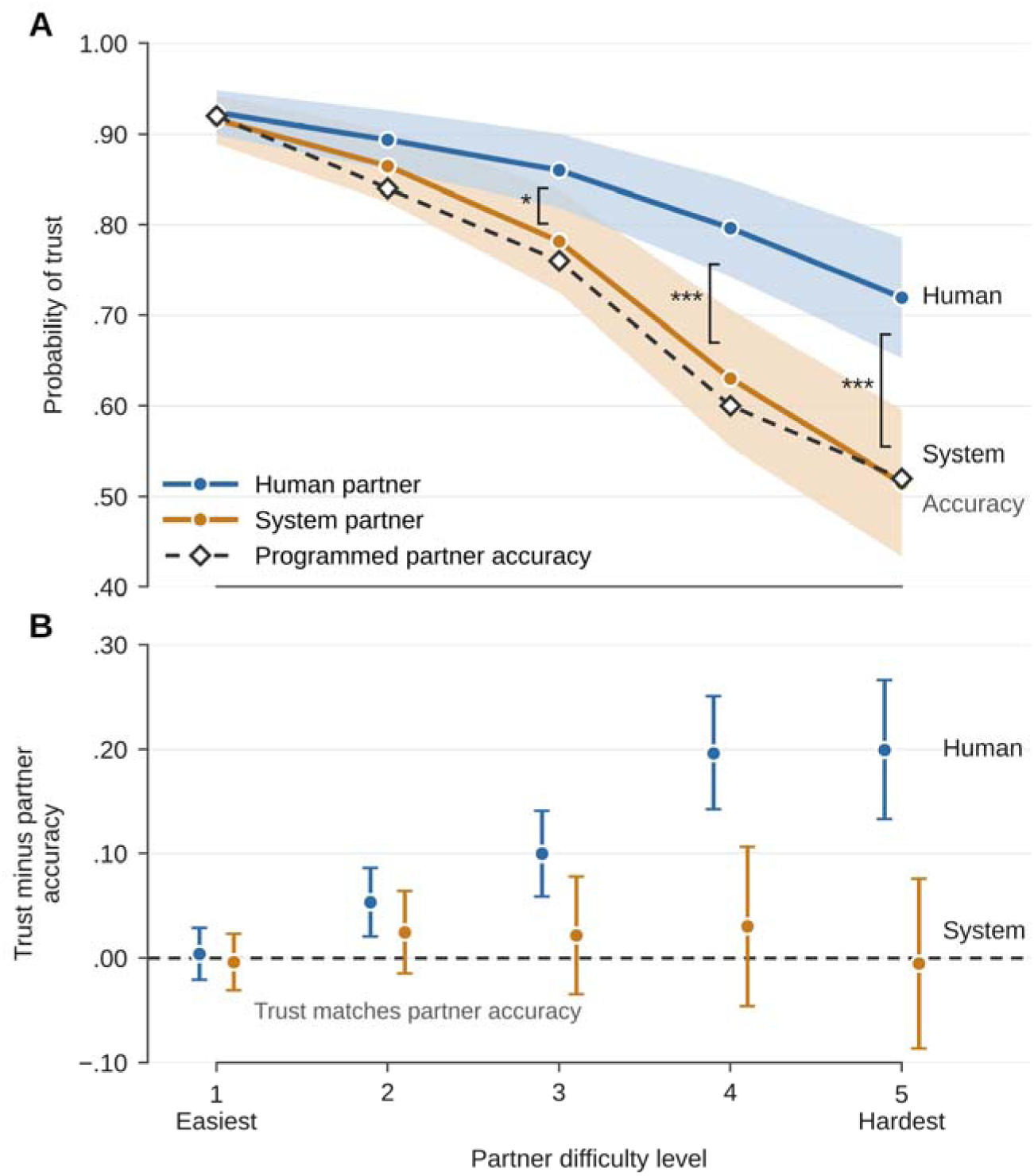
Trust in Human and System Partners Relative to Programmed Partner Accuracy. *Note.* (A) Probability of trust estimated by the binomial generalised linear mixed model for each partner condition, with shaded 95% confidence intervals, and the partner’s programmed accuracy at each difficulty level. Brackets mark significant human minus system contrasts (Table 2). (B) Trust minus programmed partner accuracy, with 95% confidence intervals. Zero indicates trust matching the partner’s accuracy. * *p* < .05. *** *p* < .001.

**Table 2.**
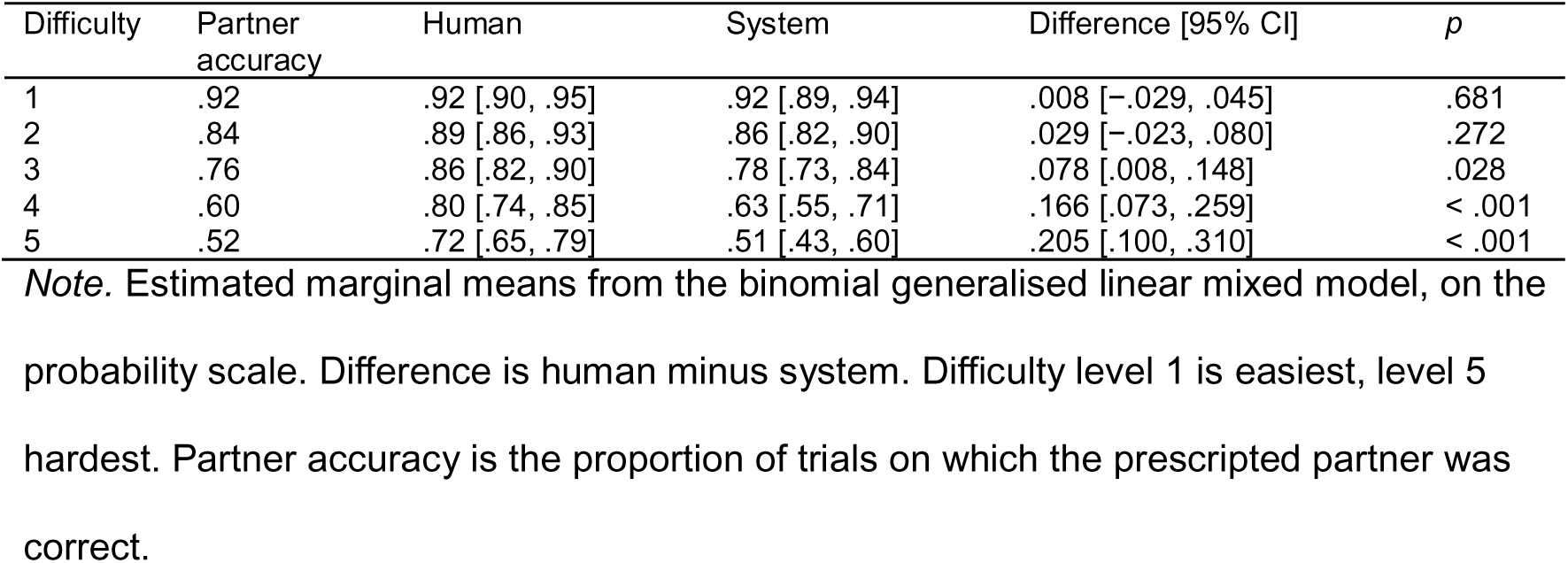
Estimated Trust Probability and Partner Contrasts by Difficulty Level.

Several participants made no distrust decisions at the lowest difficulty levels.

### Outcome Feedback Engages Performance Monitoring and Valuation Regions for Both Own and Partner Feedback

To validate task engagement, neural responses to outcome feedback were examined. For participants’ own performance, Loss > Gain elicited activation in the anterior insula and posterior medial frontal cortex (pMFC), encompassing the anterior midcingulate cortex and the supplementary and pre-supplementary motor areas, while Gain > Loss produced activation in the ventral striatum, ventromedial prefrontal cortex (vmPFC) and posterior cingulate cortex (PCC). Feedback regarding the partner’s outcome produced the same pattern for gains, in ventral striatum, vmPFC and PCC, and engaged pMFC for losses. Outcomes at partner feedback were therefore processed largely by the same performance monitoring and valuation systems as outcomes at self-performance feedback. This contrast is collapsed across partner conditions, so it characterises the processing of partner feedback in general and does not test differences between partner types (Figure 4; Table S1).

**Figure 4.**
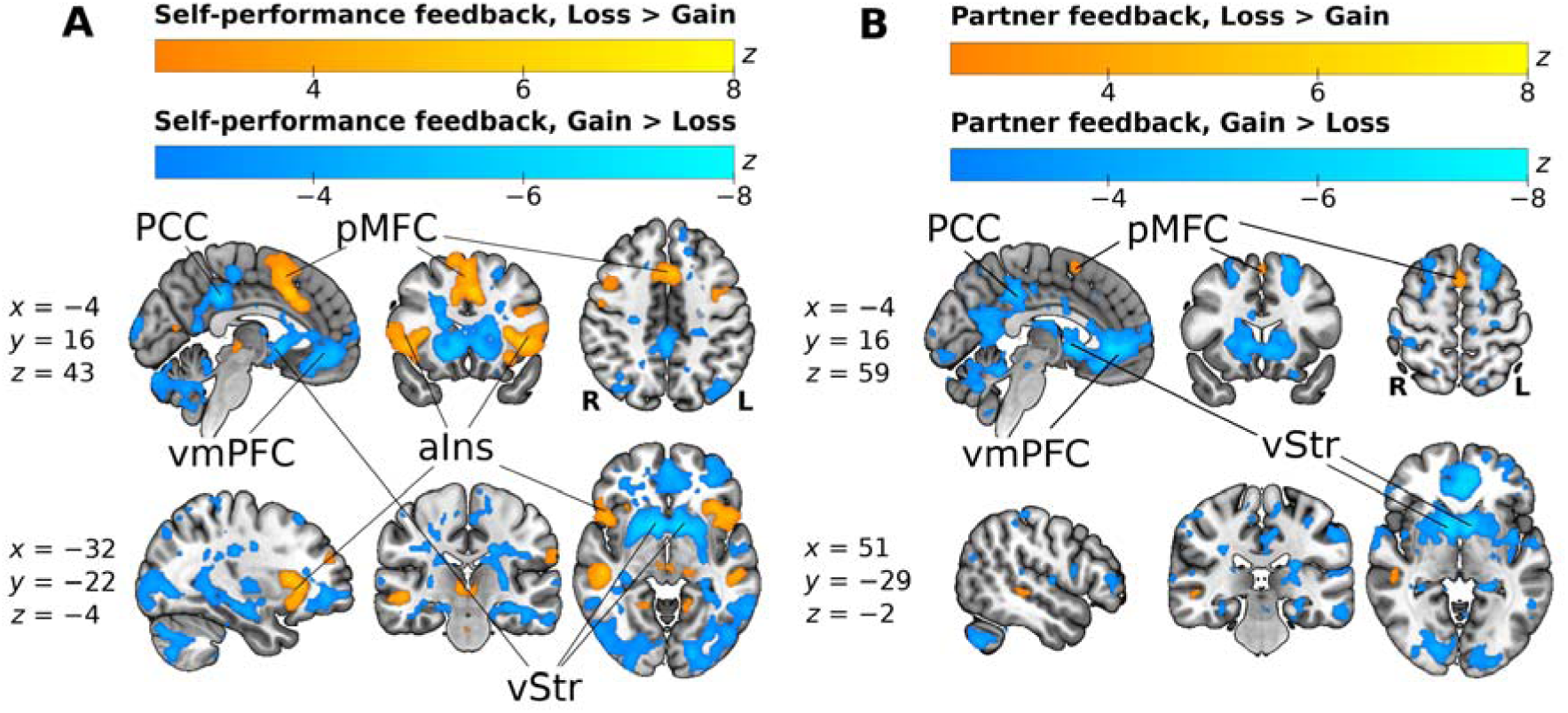
Whole-Brain Group-Level Contrasts for Outcome Feedback, Voxel-Wise FDR Corrected at q *< .05*. *Note.* Each panel shows sagittal, coronal and axial slices overlaid on the MNI152 template, with labelled regions marked on the slice where they are clearest. (A) shows feedback on the participant’s own performance and (B) shows feedback on the partner’s outcome, with Loss > Gain in warm colours and Gain > Loss in cool colours. Gains engage ventral striatum (vStr), ventromedial prefrontal cortex (vmPFC) and posterior cingulate cortex (PCC) in both, losses engage posterior medial frontal cortex (pMFC) in both, and anterior insula (aIns) responds to the participant’s own losses. Cool colours show the negative tail of the Loss > Gain contrast (negative *z*), which corresponds to Gain > Loss and is reported with positive *z* in Table S1. Coronal and axial slices are shown in radiological convention (left hemisphere on the right). Maps show only voxels surviving correction, and colour bars give *z* statistics. Slice coordinates are in MNI space (mm). Cluster statistics for every contrast are given in Table S1.

### Distrust Decisions Engage the Full Evaluative Network

Across all participants, the contrast Distrust > Trust revealed significant activation in a distributed network including bilateral anterior insula, anterior midcingulate cortex (aMCC), pre-supplementary motor area, lateral orbitofrontal cortex (lOFC), right inferior frontal gyrus (rIFG), ventral striatum, left amygdala, and right superior temporal sulcus (voxel-wise FDR corrected; Table S1). This pattern is consistent with engagement of performance monitoring, salience, and social evaluation systems during distrust decisions. Activation is shown in Figure 5; ROI coordinates are given in Table 1. The complementary contrast, Trust > Distrust, was far more limited, comprising 10,426 voxels (83.4 cm³, 11.3% of suprathreshold voxels), with clusters in ventromedial prefrontal and medial orbitofrontal cortex (peak at *x* = −6, *y* = 36, *z* = −12; peak *z* = 4.78) and bilateral posterior hippocampus. The ventromedial prefrontal cluster corresponds in location to the Gain > Loss feedback contrast reported above.

**Figure 5.**
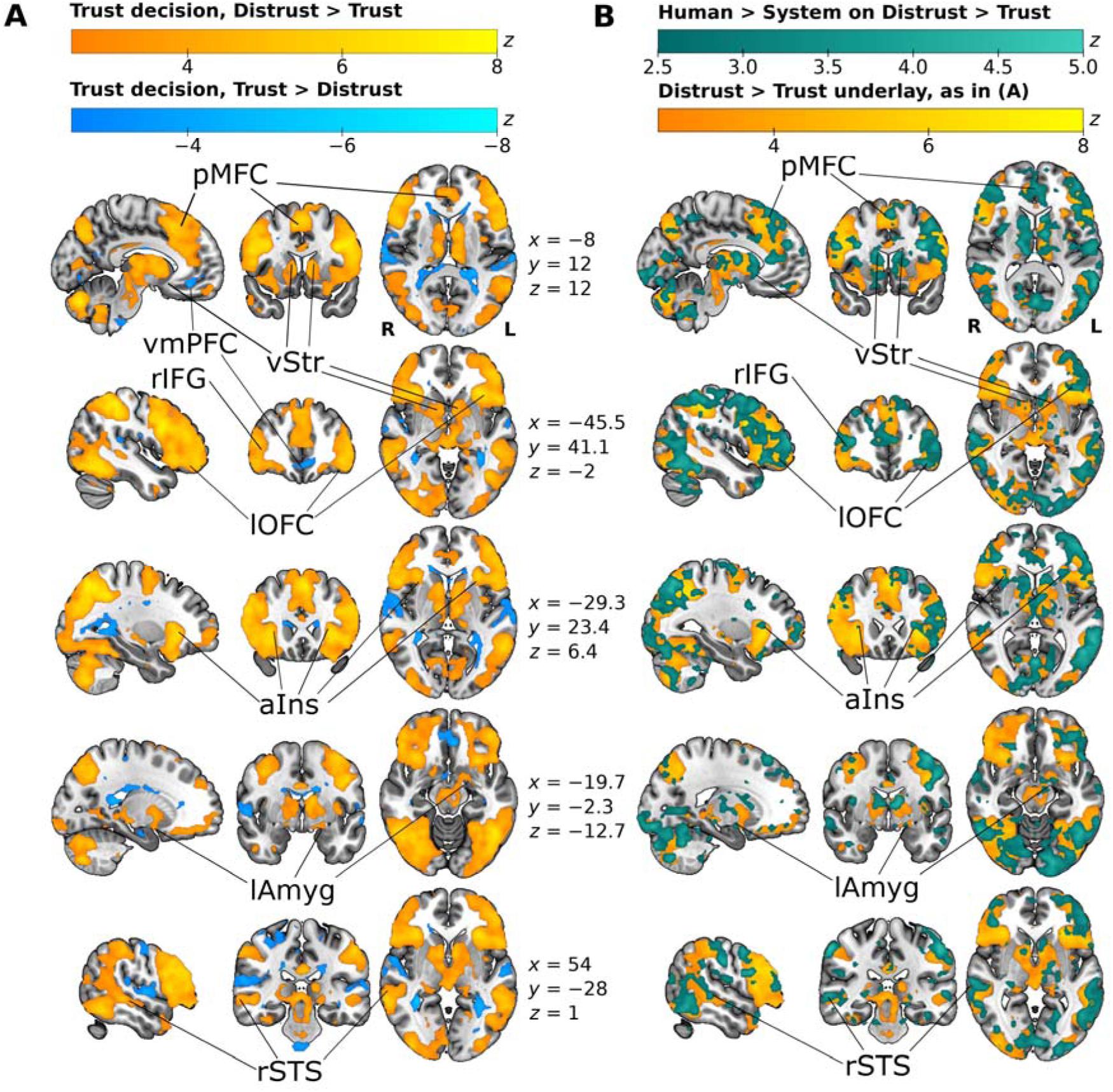
Whole-Brain Group-Level Contrasts for the Partner Trust Decision, Voxel-Wise FDR Corrected at q < .05. *Note.* Each panel shows sagittal, coronal and axial slices overlaid on the MNI152 template, with labelled regions marked on the slice where they are clearest. (A) shows the partner trust decision, with Distrust > Trust in warm colours and Trust > Distrust in cool colours. Distrust engages the distributed evaluative network, including bilateral anterior insula (aIns), posterior medial frontal cortex (pMFC), lateral orbitofrontal cortex (lOFC), right inferior frontal gyrus (rIFG), ventral striatum (vStr), left amygdala (lAmyg) and right superior temporal sulcus (rSTS). The more limited Trust > Distrust response includes ventromedial prefrontal cortex (vmPFC), in the same location as the Gain > Loss feedback response (Figure 4). Cool colours show the negative tail of the Distrust > Trust contrast (negative *z*), which corresponds to Trust > Distrust and is reported with positive *z* in Table S1. (B) shows the Human > System partner group difference on the Distrust > Trust contrast in teal, overlaid on the Distrust > Trust map from (A), with lighter teal indicating a larger difference. The partner difference engages the same distributed evaluative network as (A), with 70% of its voxels inside the Distrust > Trust map and suprathreshold voxels in all ten regions of interest. Coronal and axial slices are shown in radiological convention (left hemisphere on the right). Maps show only voxels surviving correction, and colour bars give *z* statistics. Slice coordinates between the panels are in MNI space (mm) and apply to both (A) and (B). Cluster statistics for every contrast are given in Table S1.

Direct comparison of partner groups showed greater distrust-related activation for human than for autonomous system partners (45,982 voxels, 367.9 cm³; peak in left inferior frontal gyrus at *x* = −36, *y* = 12, *z* = 24; peak *z* = 5.35), and no region showed the reverse. The effect lay predominantly within the distrust network, with 70% of its voxels inside the Distrust > Trust map and suprathreshold voxels in all ten ROIs. These results indicate that the neural architecture supporting distrust decisions is shared across partner types but exhibits increased engagement during interaction with human partners (Figure 5; Table S1).

### Trust and Distrust States Are Decodable for Both Partners and More So for Humans

Multivariate pattern analysis demonstrated that trust and distrust decisions were decodable above chance in the pooled sample (64.3%, *t*(41) = 6.90, *q* < .001, *d* = 1.06). Decoding was significant within each partner condition, at 69.8% for human partners (*t*(19) = 7.01, *q* < .001) and 59.3% for system partners (*t*(21) = 3.53, *q* = .003) and was higher for human than for system partners (difference = 10.5%, 95% CI [2.7, 18.3], *t*(39.45) = 2.72, *q* = .013, *d* = 0.84). Two aspects of this result should be distinguished. Decoding was above chance in both conditions, indicating that the neural patterns accompanying trust and distrust decisions were separable during system interaction as well as during human interaction. The difference between conditions is one of degree, with these states more discriminable during interaction with a human partner. Decoding accuracies are shown in Figure 6, full statistics are given in Table 3, and decoding against chance in Table S2.

**Figure 6.**
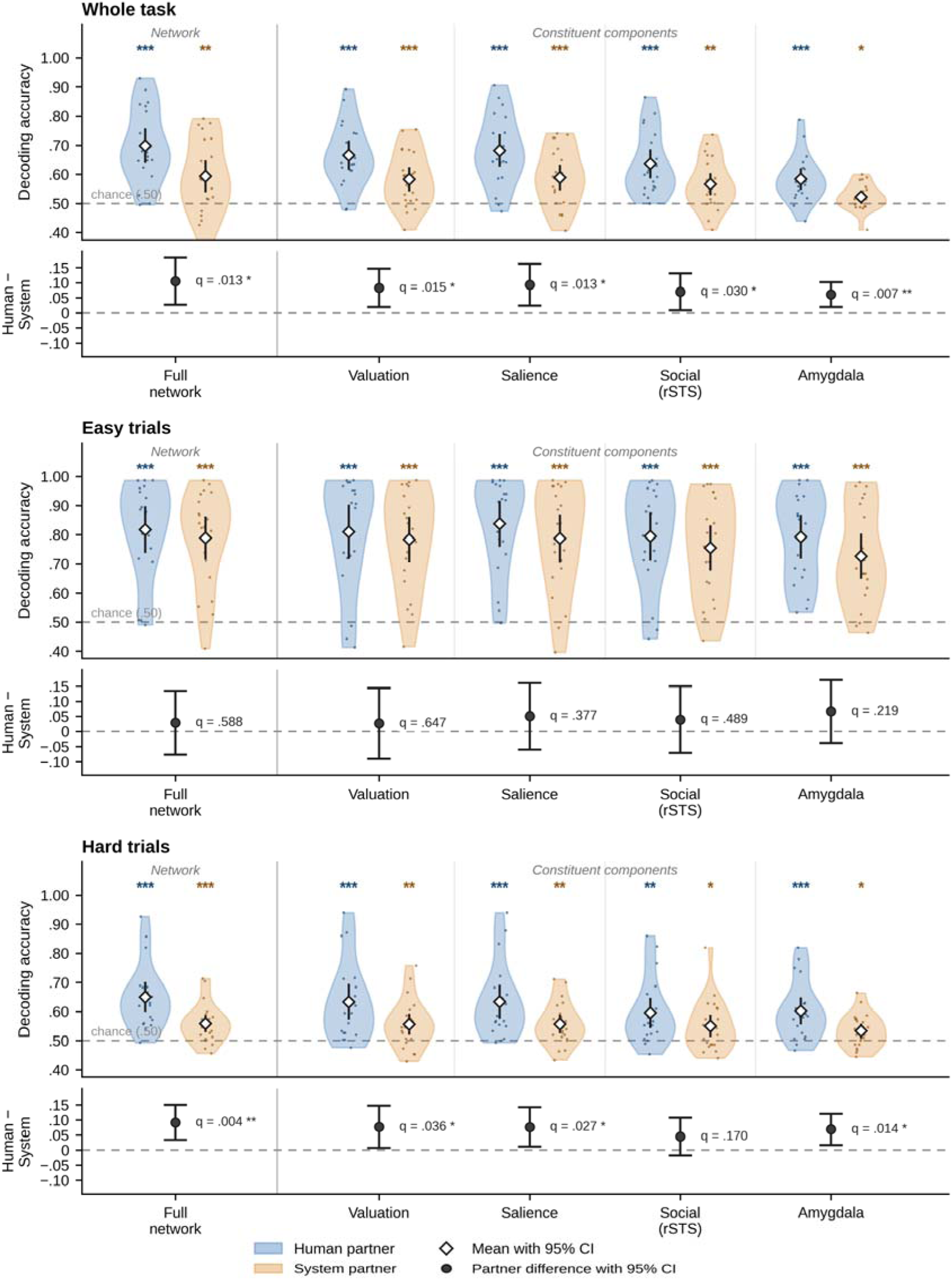
Multivariate Decoding of Trust Versus Distrust Decisions by Partner Condition. *Note.* Upper axes show within-participant cross-validated classification accuracy for each region-of-interest group; violins and points show individual participants, diamonds show the mean with 95% confidence interval, and the dashed line marks chance (.50). Stars above each violin indicate decoding above chance. Lower axes show the difference in mean accuracy between partner conditions (human minus system) with 95% confidence intervals from Welch independent-samples *t* tests. Full network features are the concatenation of the four ROI groups and are therefore not independent of them. All *q* values are Benjamini–Hochberg false discovery rates corrected across all 60 tests. Whole task *n* = 20 human, 22 system; easy *n* = 19 and 21; hard *n* = 20 and 22. * *q* < .05, ** *q* < .01, *** *q* < .001.

**Table 3.** Partner Differences in Decoding of Trust Versus Distrust Decisions.

| ROI group | Human *M* (*SD*) | System *M* (*SD*) | Difference [95% CI] | *t* | *df* | *q* | *d* |
|---|---|---|---|---|---|---|---|
| Whole task |  |  |  |  |  |  |  |
| Full network | .698 (.126) | .593 (.124) | .105 [.027, .183] | 2.72 | 39.45 | .013 | 0.84 |
| Constituent components |  |  |  |  |  |  |  |
| Valuation | .666 (.108) | .583 (.093) | .083 [.019, .146] | 2.64 | 37.82 | .015 | 0.82 |
| Salience | .682 (.120) | .589 (.099) | .093 [.024, .162] | 2.72 | 36.91 | .013 | 0.85 |
| Social (rSTS) | .636 (.108) | .566 (.086) | .070 [.008, .131] | 2.31 | 36.25 | .030 | 0.72 |
| Amygdala | .584 (.080) | .523 (.044) | .061 [.020, .102] | 3.03 | 28.85 | .007 | 0.96 |
| Easy trials |  |  |  |  |  |  |  |
| Full network | .818 (.166) | .788 (.163) | .029 [−.076, .135] | 0.56 | 37.41 | .588 | 0.18 |
| Constituent components |  |  |  |  |  |  |  |
| Valuation | .810 (.193) | .783 (.171) | .027 [−.091, .144] | 0.46 | 36.20 | .647 | 0.15 |
| Salience | .837 (.163) | .787 (.181) | .051 [−.060, .161] | 0.93 | 38.00 | .377 | 0.29 |
| Social (rSTS) | .794 (.173) | .755 (.171) | .040 [−.071, .150] | 0.73 | 37.50 | .489 | 0.23 |
| Amygdala | .793 (.156) | .726 (.171) | .067 [−.038, .171] | 1.29 | 38.00 | .219 | 0.41 |
| Hard trials |  |  |  |  |  |  |  |
| Full network | .650 (.110) | .558 (.066) | .092 [.034, .150] | 3.25 | 30.55 | .004 | 1.03 |
| Constituent components |  |  |  |  |  |  |  |
| Valuation | .633 (.132) | .556 (.082) | .077 [.007, .147] | 2.25 | 31.16 | .036 | 0.71 |
| Salience | .634 (.125) | .557 (.072) | .077 [.011, .142] | 2.40 | 29.82 | .027 | 0.76 |
| Social (rSTS) | .595 (.111) | .550 (.086) | .045 [−.018, .108] | 1.45 | 35.76 | .170 | 0.45 |
| Amygdala | .602 (.100) | .533 (.056) | .069 [.017, .120] | 2.71 | 29.31 | .014 | 0.86 |

*Note.* Values are within-participant cross-validated classification accuracies for trust versus distrust decisions (LinearSVC, five-fold stratified cross-validation); chance = .50. Difference = human minus system, with 95% confidence interval. Comparisons are Welch independent-samples *t* tests, as partner condition was manipulated between subjects. *q* = Benjamini–Hochberg false discovery rate, corrected across all 60 tests reported in Table 3 and Table S2. *d* = Cohen’s *d* for independent samples. Full network features are the concatenation of the four ROI groups and are therefore not independent of them. Whole task *n* = 20 human, 22 system; easy *n* = 19 and 21; hard *n* = 20 and 22. Two participants in the human condition were excluded from all MVPA analyses (see Methods). Easy = difficulty levels 1 and 2; hard = levels 4 and 5; level 3 was excluded to maximise contrast between difficulty extremes. Participants contributing trials of only one decision type within a bin were excluded from that bin. Decoding exceeded chance in every cell (Table S2).

### Attenuation for System Partners Is Present in Every Network Component

ROI-based analyses confirmed above-chance decoding across valuation, salience, social, and amygdala networks, with significant human > system differences in each (Table 3). Accuracies for human and system partners respectively were 66.6% and 58.3% in valuation regions (*q* = .015, *d* = 0.82), 68.2% and 58.9% in salience regions (*q* = .013, *d* = 0.85), 63.6% and 56.6% in the social ROI (*q* = .030, *d* = 0.72), and 58.4% and 52.3% in the amygdala (*q* = .007, *d* = 0.96). Every component carried decision-relevant information during system interaction, and every component carried less of it than during human interaction. Attenuation was therefore present throughout the evaluative architecture, and the partner differences were large (*d* = 0.72 to 0.96). The amygdala showed the smallest raw difference (6.1%) with the largest standardised effect of the four components (*d* = 0.96).

### Partner Differences in Decoding Emerge When Partner Accuracy Is in Doubt

Difficulty-stratified analyses were restricted to participants contributing decisions of both types within a bin (easy *n* = 19 human, 21 system). Under easy conditions, decoding accuracy remained significantly above chance across all networks, but no differences were observed between partner groups (full network 81.8% human and 78.8% system, difference = 2.9%, 95% CI [−7.6, 13.5], *q* = .588; valuation *q* = .647; salience *q* = .377; social *q* = .489; amygdala *q* = .219).

Under hard conditions, decoding remained above chance in all networks and in both partner conditions, and partner differences emerged. At the network level across all ROIs, decoding accuracy was higher in the human group than in the system group (65.0% and 55.8%, difference = 9.2%, 95% CI [3.4, 15.0], *t*(30.55) = 3.25, *q* = .004, *d* = 1.03). Among individual ROIs, the difference was significant in valuation regions (63.3% and 55.6%, *q* = .036, *d* = 0.71), salience regions (63.4% and 55.7%, *q* = .027, *d* = 0.76) and the amygdala (60.2% and 53.3%, *q* = .014, *d* = 0.86). The difference in the social ROI (59.5% and 55.0%, *q* = .170) did not reach the corrected threshold. The whole-network effect was larger under hard trials (*d* = 1.03) than across the task as a whole (*d* = 0.84).

### Trust and Distrust Decisions Recruit the Same Coupled Network for Both Partner Types

Within-group coupling was significant for all 45 region pairs in every cell of the design, in hard trust and in hard distrust trials and in both partner groups (all *q* < .05). The ten regions therefore operated as a comprehensively coupled system during both decision types, irrespective of partner type (Figure 7; Table 4). Median coupling was 0.24 for human trust, 0.25 for system trust, 0.28 for human distrust and 0.22 for system distrust. Coupling was organised by ROI membership and was not distributed evenly. Within hard distrust trials, mean coupling among the four salience regions was 0.517 in the human group and 0.456 in the system group, higher than among the four valuation regions (0.256 and 0.219) or between regions of different networks (0.287 and 0.242). Full within-group statistics are given in Tables S3 and S4.

**Figure 7.**
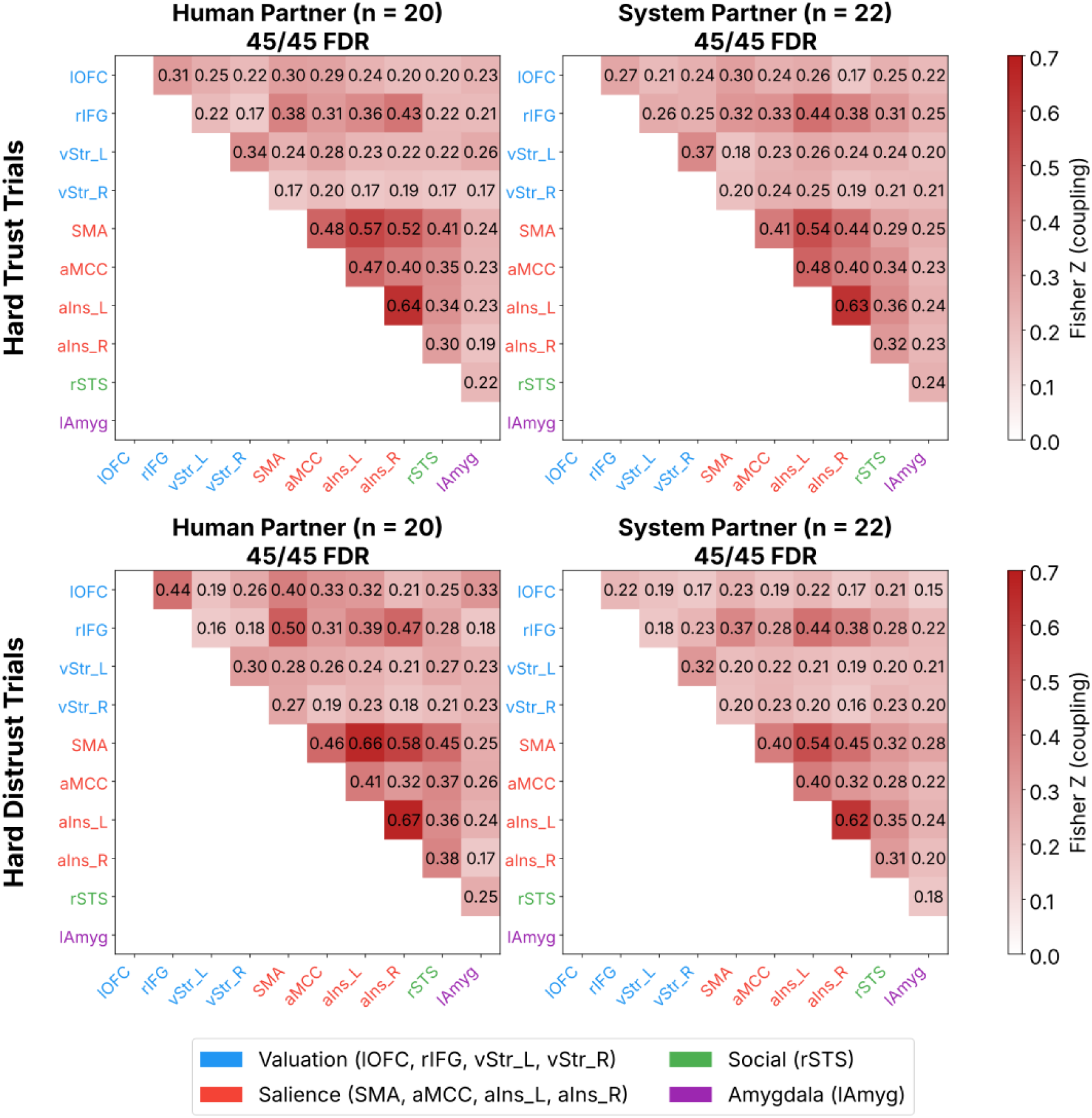
Beta-Series Functional Connectivity Between the Ten Regions of Interest on Hard Trials. *Note.* Rows show hard trust trials (upper) and hard distrust trials (lower). Columns show mean Fisher *z* coupling for the human partner group (*n* = 20) and the system partner group (*n* = 22). In every panel all 45 region pairs survived FDR correction against zero within group, as indicated above each panel. Region labels are coloured by network assignment.

**Table 4.** Beta-Series Functional Connectivity, Within- and Between-Group Coupling by Decision Type.

| Decision type | Group | *n* | Pairs sig. (of 45) | *Mdn z* | Range | Salience | Valuation | Cross-network |
|---|---|---|---|---|---|---|---|---|
| Hard trust | Human | 20 | 45 | 0.237 | 0.169 to 0.635 | 0.513 | 0.251 | 0.254 |
|  | System | 22 | 45 | 0.247 | 0.175 to 0.631 | 0.483 | 0.267 | 0.261 |
|  | Human − System |  | 0 | −0.016 | −0.097 to +0.118 |  |  |  |
| Hard distrust | Human | 20 | 45 | 0.275 | 0.161 to 0.673 | 0.517 | 0.256 | 0.287 |
|  | System | 22 | 45 | 0.224 | 0.152 to 0.621 | 0.456 | 0.219 | 0.242 |
|  | Human − System |  | 0 | +0.034 | −0.046 to +0.213 |  |  |  |

*Note. Mdn z* is the median Fisher *z*-transformed correlation across the 45 ROI pairs. Pairs sig. = number of the 45 ROI pairs significant after false discovery rate correction (Benjamini–Hochberg, *q* < .05); within-group rows test coupling against zero (one-sample *t* tests), and Human − System rows test the group difference (Welch *t* tests). For Human − System rows, the *Mdn z* column reports the median difference and Range reports the minimum and maximum difference across the 45 pairs. Salience, Valuation, and Cross-network columns give mean coupling within the four salience regions, within the four valuation regions, and between regions of different networks, respectively; these are descriptive and were not tested inferentially.

No between-group difference survived correction in either decision type (minimum *q* = .944 in hard trust, .714 in hard distrust). Coupling also did not differ between trust and distrust decisions in either partner group, with no pair surviving correction. Full statistics are given in Tables S5 to S7, and the complete comparison grid in Supplementary Figure S2.

## Discussion

This study asked whether trust decisions towards an autonomous system engage the evaluative network that supports trust between people, and how that engagement depends on uncertainty. The two partners performed identically, yet trust was withdrawn more readily from the system partner once partner errors became frequent, tracking its actual accuracy at every difficulty level, whereas trust in the human partner persisted above the same accuracy from level 2 onward. The regions specified in advance from the performance monitoring framework were comprehensively coupled on hard trials during trust and distrust decisions in both partner conditions. Interaction with an autonomous system did not disengage this network but attenuated it, in activation and in the discriminability of trust and distrust states, in every component examined, whereas our earlier passive monitoring study found the P3a to a system partner’s errors absent altogether (Rogers et al., 2026). The decoding difference was expressed under high difficulty, where behaviour also separated the partners. These findings suggest that trust in an autonomous system recruits the same evaluative architecture as trust in a person, but with attenuated engagement accompanied by trust that more closely tracks the partner’s actual reliability.

### Distrust as Prospective Performance Monitoring

Trust was the default in this task. At low difficulty several participants never distrusted, and distrust rose only as partner performance declined. We therefore treat a decision to distrust as a departure from reliance, made on the judgement that the partner is about to fail, and as the prospective counterpart of error detection. Performance monitoring theory predicts that such a judgement recruits posterior medial frontal cortex, which responds to decision uncertainty and unfavourable outcomes as well as to errors (Ridderinkhof et al., 2004; Ullsperger et al., 2014a), and anterior insula, whose activity tracks risk and risk prediction error (Preuschoff et al., 2008). Consistent with this, posterior medial frontal cortex responded to participants’ own losses, to their losses at partner feedback and to the decision to distrust, and anterior insula to own losses and to distrust. Trust carried its own, more limited response, which included ventromedial prefrontal cortex, a core region of the valuation system (Bartra et al., 2013), at the location that responded to gains at feedback, consistent with valuation processes engaged when anticipating partner success, as distrusting engaged performance monitoring when anticipating partner failure.

Performance monitoring can extend to a partner’s anticipated failure because it is engaged by another agent’s outcomes as well as one’s own. Observed errors recruit posterior medial frontal cortex, anterior insula and posterior superior temporal sulcus at least as strongly as self-generated errors (Monfardini et al., 2013), and medial frontal neurons in macaques respond selectively to a partner’s errors, in populations distinct from those signalling the animal’s own errors but within the same region, so fMRI cannot separate them (Yoshida et al., 2012). This indicates that the system is agent-general with respect to the source of the error. If so, the network should be recruited for both partners, and a partner effect should lie in the valuation, social and affective input signals it receives.

### A Coupled Evaluative Network

The activation and decoding analyses show that each component carries decision-relevant information, but not that the components operate together. On hard trials, coupling was significant for all 45 region pairs in both partner conditions and in both decision types, so valuation, salience, social-perceptual and affective regions covary trial by trial whenever a judgement about the partner is made, not only when the partner is doubted. This is the empirical basis for treating trust in autonomous systems as a network phenomenon rather than a collection of regional effects. The constituent regions are also implicated in interpersonal trust and social evaluation (Bellucci et al., 2017; Isik et al., 2017; Rosenberger et al., 2019). The coupling result also tests directly, within a network specified in advance, the proposal that trust depends on interactions among large-scale neural systems and not on regional activity alone (Krueger & Meyer-Lindenberg, 2019), and supports it for both partner types.

Coupling was not uniform. Coupling among the four salience regions was roughly twice that among the four valuation regions in both partner groups, in hard distrust and in hard trust trials, consistent with the strong intrinsic coupling that defines the salience network (Uddin, 2015). This graded structure is more consistent with organisation within the evaluative system than with a single task-evoked signal shared equally by every region.

The three neural measures are dissociable, as activation and decoding both differ between partner conditions, whereas no differences in coupling between partners or between decision types survived correction on hard trials. This suggests that the architecture engaged by a trust judgement is shared across partner type and decision type, while the strength of engagement and the separability of decision states within it vary. Connectivity differences have been reported between advice from a machine and from a person (Goodyear et al., 2016, 2017), and network-level comparison of human-robot and human-human interaction has found distinct profiles across perceptual and language systems (Hogenhuis & Hortensius, 2022). Because those studies define the network from the data within each condition, a difference between conditions can reflect either a different architecture or the same architecture engaged to a different degree. Specifying the network in advance separates the two.

### Attenuation Rather Than Absence

Attenuation appears in two different measures as distrust-related activation was greater for human than for system partners, with the difference present in all ten regions, while trust and distrust states were more separable for human partners in every component. These index different things, the overall response and the separability of the two decision states, and they agree in direction.

In our earlier EEG study, under passive monitoring, errors by an autonomous system partner elicited no P3a while errors by a human partner did, although the feedback-related negativity and P3b to those errors were preserved and behaviour did not differ between partners (Rogers et al., 2026). Basic error detection and outcome evaluation thus transferred to the system partner, while the frontal attentional engagement indexed by the P3a (Polich, 2007) did not, and the partner difference was evident in the brain alone. We left open whether technology-related cognitive economization (TRCE) would extend from passive observation to active decisions about a system’s trustworthiness, where we suggested frontal engagement might be recovered. The present findings suggest that TRCE does extend to active decisions, in a different form, because the requirement to decide appears to impose a floor of engagement, so that TRCE is expressed as attenuation rather than absence. In contrast to our earlier work, the partner difference here was also expressed in behaviour. Trust was withdrawn more readily from the system partner as its reliability declined, so a difference in neural engagement that passive monitoring left without a behavioural counterpart was accompanied by one once the partner had to be evaluated in order to decide. Establishing when differences in neural processing translate into differences in behaviour is an important direction for future research.

In our view, attribution of agency to the partner is a plausible cause of this attenuation, since social responses to artificial agents depend on whether the observer attributes intentions to them (Wiese et al., 2017), and, it has been argued, more on that attribution than on the agent’s features (Hortensius & Cross, 2018). A human partner’s mistakes may be more readily attributed to circumstance, effort, or a bad run, whereas a system’s errors may be interpreted more directly as evidence of its reliability. Consequently, social-perceptual and affective components may contribute more strongly when the partner is credited with acting instead of simply producing outputs.

It is important to note that attenuation is visible only against a human reference. In isolation, the system condition, with above-chance decoding in every component, could be read as fully engaged processing. Much of the neuroscience of human-automation interaction examines responses to automation without a matched human partner (Hopko & Mehta, 2021), and so cannot distinguish typical from diminished processing.

### Difficulty as a Contextual Modulator

Decoding separated the partners under high difficulty, at the whole-network level and in the valuation, salience and amygdala components, but not under easy conditions. Behaviour, an index independent of the neural data, followed the same course, matched while the partner performed well and separating as partner errors became frequent. This is what a performance monitoring account predicts. Monitoring is engaged in proportion to uncertainty (Grinband et al., 2006; Ridderinkhof et al., 2004), and anterior cingulate cortex tracks how far information supplied by another agent should guide choice (Behrens et al., 2008), so a partner effect should be expressed most strongly where the partner’s accuracy is in doubt and the decision must rest on what has been represented about that partner. When the partner difference appears rests on the behavioural model, in which trust diverged from level 3 onward. The easy condition yields too few distrust trials to resolve a partner difference of the magnitude observed under hard conditions, and is therefore not interpreted as a null. Lowering partner accuracy in the easiest conditions may yield more distrust decisions there, and with them a more sensitive test of the partner difference where the partner is largely reliable.

### Social-Affective Evaluation of Artificial Agents

Whether artificial agents engage the systems that evaluate other people is a long-standing question in social neuroscience, and the evidence supports two readings. Some responses are categorical, for example when every opponent’s play was identical and only the believed opponent differed, medial prefrontal cortex and right temporoparietal junction responded to a human opponent and not to a robot or a computer (Chaminade et al., 2012). Others however, are graded, with studies showing right temporoparietal junction and medial frontal activity rose linearly with a partner’s human-likeness, from computer through two robots to a human (Krach et al., 2008), and trust-related EEG modulation is stronger for more human-like agents (Jung et al., 2019). Partner studies report weaker anterior insula responses to unfair offers from a computer than from a human (Sanfey et al., 2003), weaker striatal responses to cooperation with a computer partner (Rilling et al., 2002) and weaker amygdala responses to technological partners (Rauchbauer et al., 2019), and amygdala activity predicts the rejection of artificial partners (Rosenthal-von der Pütten et al., 2019).

In the present data, the whole-task analysis shows attenuation in every component, including the social-perceptual and affective components. Our earlier interpretation of the absent P3a as TRCE, a reduced allocation of social and affective resources, could not be tested with EEG, which cannot separate social from affective processing. The present data allow that interpretation to be examined region by region, and in our view support it asymmetrically. The amygdala, which assigns social-affective value to a partner’s anticipated action, showed the largest standardised partner effect of the four components, *d* = 0.96 across the task and 0.86 under high difficulty, despite the smallest raw difference in decoding accuracy across the task, converging with evidence that basolateral amygdala damage abolishes adaptive updating of investment in a repeated trust game (Rosenberger et al., 2019). The salience components, in which anterior insula and aMCC register the aversive and motivational weight of an uncertain outcome (Craig, 2009; Shackman et al., 2011), were attenuated alongside the amygdala. These frontal regions overlap the circuitry thought to generate the P3a (Polich, 2007), so the P3a absent in our earlier study and the salience attenuation here may reflect one system at two phases. The social-perceptual attenuation is established across the task but falls below threshold under high difficulty.

Set against this literature, we suggest the present results favour the graded reading for the components examined. The system partner engaged the social-perceptual and affective components, but less than the human partner did, so the difference lay in how strongly a partner was evaluated rather than in whether it was evaluated socially. Because the two partners performed identically, that difference arose from what participants believed about the partner, which is consistent with the attribution account proposed above.

### Trust Calibration and Implications for Trust in Automation

Trust in automation has been studied as a behavioural construct and as a neural signal, with little connecting the two. Behaviourally, trust in automation is less resilient to violation than interpersonal trust (de Visser et al., 2016; Madhavan & Wiegmann, 2007), and situational factors including task difficulty modulate it (Hoff & Bashir, 2015). Recent reviews treat trust in automation and in artificial intelligence as one problem (Chen et al., 2025).

Neural correlates have been sought one signal or one region at a time, including an error-monitoring potential tracking trust in an algorithm (de Visser et al., 2018) and prefrontal activation under robot unreliability (Hopko et al., 2024), alongside the partner effects in social and affective regions reviewed above. These correlates vary widely across paradigms and technology domains (Hopko & Mehta, 2021), and although they point in the same direction they have remained a list of findings rather than an account, with the call to bring decision neuroscience to the problem not yet taken up at the network level (Drnec et al., 2016).

The present study connects these levels. Read against a network specified in advance, the correlates reported across these studies reflect graded engagement of one architecture, and the readiness with which trust in automation is withdrawn, documented behaviourally as algorithm aversion (Dietvorst et al., 2015), may have a mechanism. We suggest that a thinner evaluative representation, evident in the reduced discriminability of trust and distrust states for the system partner, may provide less beyond the error rate to sustain trust when the partner’s accuracy becomes uncertain.

Lee and See (2004) defined appropriate reliance as trust calibrated to a partner’s capability, and distinguished analytic, analogical and affective processes underlying trust. Read as an estimate of how often the partner would be correct, trust in the system partner matched its actual accuracy at every level, while trust in the human partner exceeded it from level 2 onward. The more readily withdrawn trust was also the better calibrated. Trust in the system partner showed neither of the miscalibrations the behavioural literature describes for automation, neither exceeding its reliability, as expectations of machine perfection would lead one to expect (Dzindolet et al., 2003), nor falling below it, as algorithm aversion would suggest (Dietvorst et al., 2015). It was trust in the human partner that departed from reliability, held above a reliability identical to the system’s. Against a human benchmark, trust calibrated to a system’s reliability can therefore appear as aversion. The social-perceptual and affective components that differed between partners may contribute to sustaining trust in a human partner above its demonstrated reliability. This social-affective support for trust in a human partner may also account for the greater distrust-related engagement with that partner, since each decision to distrust a person departed further from it.

Automation-related complacency has been characterised in attentional terms, as reduced monitoring of a system presumed reliable (Parasuraman & Manzey, 2010; Parasuraman & Riley, 1997). The present results indicate what is reduced. Task engagement was not absent, since decoding remained above chance in every component and the network remained functionally coupled for system partners on hard trials. The regional differences are consistent with reduced social-affective engagement during evaluation, and the absent P3a in our earlier study (Rogers et al., 2026) may reflect a related selective reduction rather than general inattention to the task. Whether this economization is a cost therefore depends on what the task demands. Under passive monitoring, a system’s errors are detected and tracked, but without the frontal attentional engagement indexed by the P3a, the condition in which complacency develops. Under a forced decision, the same economization coincided with trust in the system that stayed with its record. This predicts that requiring explicit judgements of an automated partner at points in a task would retain the calibration seen here while countering the complacency of passive monitoring.

## Conclusion

Trust in an autonomous system engages the evaluative network that supports trust between people, and its components remain coupled as a functioning system whether the partner is trusted or distrusted. What a system partner changes is the strength of that engagement, lower in activation and in how separable trust and distrust states are, in every component, including the social-perceptual and affective ones, and the difference opens as the partner’s accuracy comes into doubt. Our earlier evidence showed that partner attribution shapes outcome evaluation, and the present findings extend that to the decision phase, where TRCE appears as a less social and less affective evaluation of the partner, with no withdrawal from the task. Its cost depends on the task at hand, as under passive monitoring it left a system’s errors detected and tracked, but without frontal attentional engagement. Whereas, under a forced decision, trust was withdrawn more readily from the system partner, but in line with its reliability, while trust in the human partner was held above it. These findings give Lee and See’s neurological level an empirical footing two decades after they named it, and they make two predictions. If attribution of agency drives the attenuation, it should follow the human-likeness gradient described above, so a language-model partner, fluent and readily described in mental terms, should be attenuated less than the system partner here, and trust in it may then exceed its reliability, as trust in the human partner did. The anticipatory phase of trust interaction also remains uncharacterised. If economization reflects reduced allocation of resources to a non-human partner, it should be present before the partner’s outcome is known, as the attenuation at the trust decision here already suggests, and most clearly under passive monitoring, where no decision imposes a floor of engagement. Reduced anticipation would then account for the absent P3a at outcome, placing the earliest partner-dependent divergence before feedback arrives.

## Declarations

## Open Practices Statement

The data for this study are available from the corresponding author on reasonable request. The study was not preregistered.

## Supplementary Materials

### Supplementary Methods

Two participants were excluded from the multivariate pattern and beta-series connectivity analyses, for separate reasons related to their single-trial (least squares separate) beta estimates. Sub-01 was excluded because of a stimulus presentation confound. For this participant the partner difficulty blocks ran to approximately 75 trials rather than the intended 25, and continued across run boundaries. Run 1 contained two blocks, run 2 contained three, and run 3 contained two, with two blocks divided across consecutive runs. Every other participant received five blocks of approximately 25 trials within each run, so that each block type recurred in every run. Single-trial estimates are computed within run, with a nuisance regressor collapsing the remaining decision trials of that run. For this participant that regressor drew on a smaller and differently constituted set of blocks than for any other participant, so the resulting trial-wise estimates do not rest on a comparable model.

Sub-35 was excluded because the single-trial model failed to converge, returning beta estimates several orders of magnitude outside the range observed across the remainder of the sample and therefore numerically invalid. This failure was specific to the single-trial model. The conventional first-level GLM converged normally for this participant, with contrast z-maps comparable in distribution to the rest of the sample (*SD* = 1.05, range −4.6 to +4.6, no non-finite values). Both participants were retained in the univariate GLM analyses. Each event is modelled with its own onset and duration, and the reported contrasts collapse across stimulus blocks, so neither the non-standard block structure nor the single-trial convergence failure affects those estimates. Multivariate and connectivity analyses therefore included 42 participants (human *n* = 20, system *n* = 22), while all 44 participants contributed to the behavioural and univariate GLM analyses.

**Figure S1.**
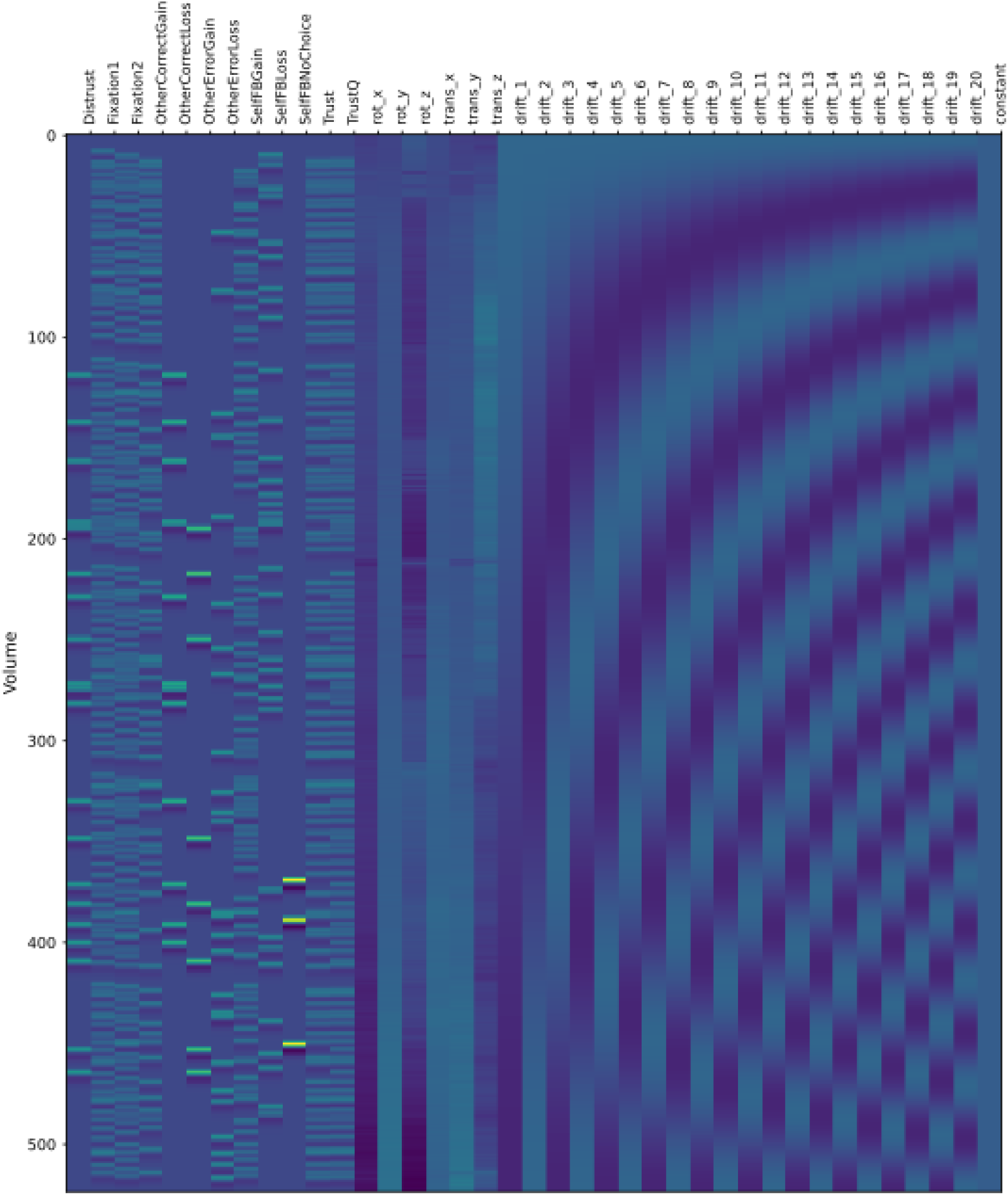
First-Level Design Matrix for a Representative Participant (sub-24, Run 1) *Note.* Columns show the twelve task event regressors convolved with the Glover canonical haemodynamic response function, the six rigid-body motion parameters, twenty cosine drift terms and the constant. Rows are volumes in acquisition order. All three runs were modelled simultaneously within a single first-level model, with a separate design matrix per run.

**Figure S2.**
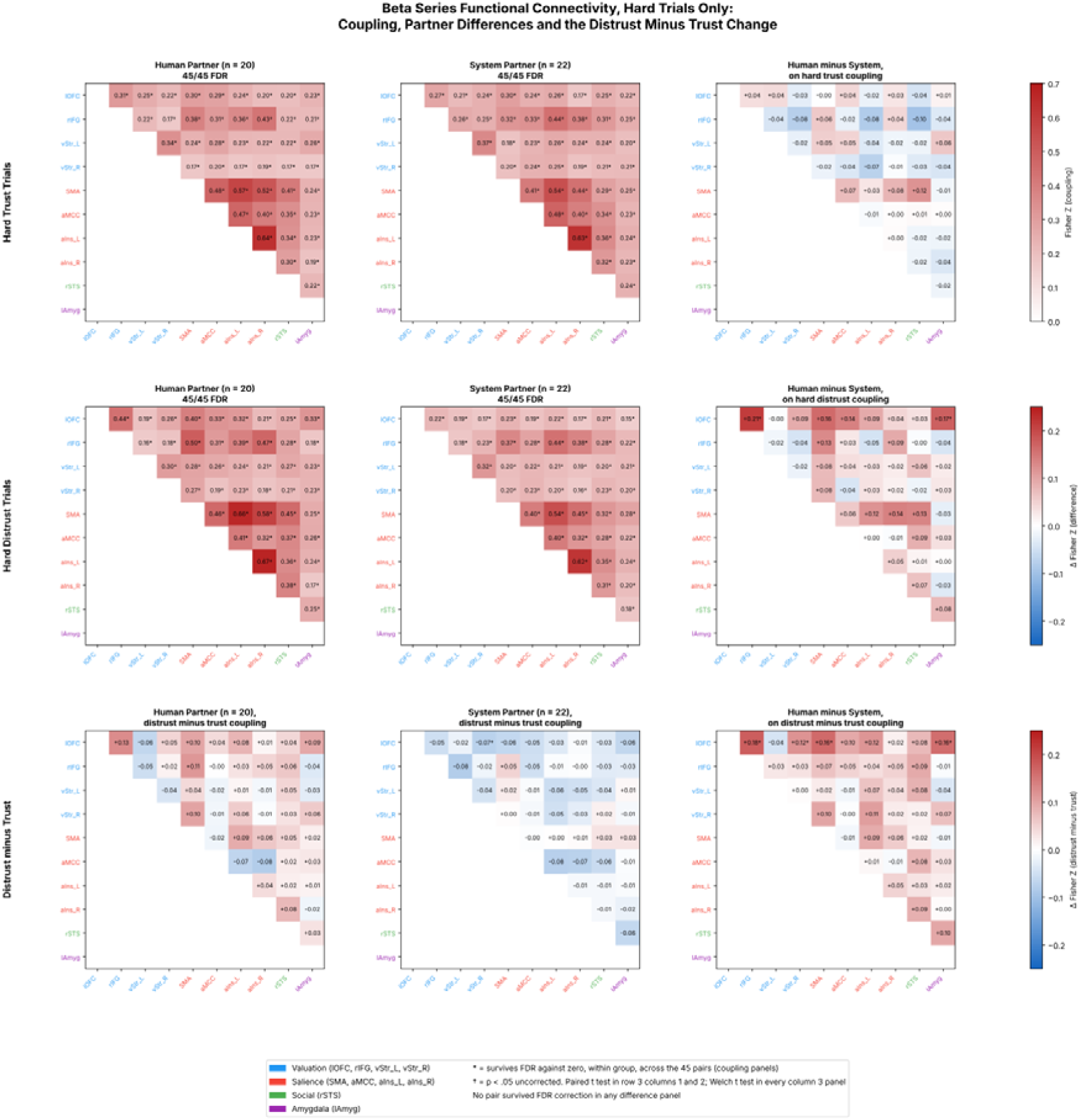
Beta-Series Functional Connectivity on Hard Trials, Showing Coupling, Partner Differences and the Distrust Minus Trust Change. *Note.* Rows show hard trust trials (upper), hard distrust trials (middle) and the distrust minus trust change (lower). Columns show the human partner group (*n* = 20), the system partner group (*n* = 22) and the human minus system difference. Asterisks in the coupling panels mark pairs surviving FDR correction against zero within group across the 45 pairs. Daggers mark differences significant at *p* < .05 uncorrected, from paired *t* tests in the first two panels of the lower row and Welch *t* tests in every panel of the third column. No pair survived FDR correction in any difference panel.

**Table S1.** Whole-Brain Clusters for the Feedback and Trust Decision Contrasts.

| Region | *x* | *y* | *z* | Peak *z* | Extent (mm³) |
|---|---|---|---|---|---|
| *Own performance feedback, Loss > Gain (8,578 voxels; minimum surviving z = 2.58)* |  |  |  |  |  |
| Left frontal operculum cortex | −42 | 18 | 0 | 5.99 | 14,408 |
| Left paracingulate gyrus | −4 | 16 | 40 | 5.90 | 15,960 |
| Right middle temporal gyrus, posterior division | 50 | −26 | −6 | 5.34 | 3,920 |
| Right frontal operculum cortex | 44 | 14 | 0 | 5.23 | 8,168 |
| Right inferior temporal gyrus, anterior division | 48 | 2 | −36 | 4.92 | 2,616 |
| Right angular gyrus | 52 | −48 | 30 | 4.67 | 3,904 |
| Right thalamus | 8 | −18 | 2 | 4.60 | 1,336 |
| Left angular gyrus | −62 | −56 | 24 | 4.52 | 2,072 |
| Right lingual gyrus | 28 | −54 | 2 | 4.43 | 4,632 |
| Left middle temporal gyrus, posterior division | −54 | −30 | −4 | 4.40 | 896 |
| Left temporal pole | −48 | 2 | −36 | 3.91 | 600 |
| Left postcentral gyrus | −66 | −22 | 28 | 3.79 | 896 |
| Right frontal pole | 24 | 52 | 26 | 3.77 | 832 |
| Left intracalcarine cortex | −16 | −68 | 6 | 3.76 | 2,560 |
| Left middle frontal gyrus | −40 | 0 | 48 | 3.76 | 1,840 |
| Right middle frontal gyrus | 46 | 30 | 32 | 3.58 | 480 |
| Left frontal pole | −28 | 50 | 22 | 3.47 | 1,328 |
| Right middle frontal gyrus | 46 | 8 | 44 | 3.43 | 856 |
| *Own performance feedback, Gain > Loss (36,115 voxels; minimum surviving z = 2.58)* |  |  |  |  |  |
| Right putamen | 14 | 8 | −8 | 7.86 | 246,472 |
| Left accumbens | −12 | 6 | −8 | 7.81 |  |
| Left inferior temporal gyrus, posterior division | −54 | −42 | −12 | 6.97 |  |
| Right lateral occipital cortex, inferior division | 44 | −76 | −32 | 6.19 |  |
| Left cingulate gyrus, posterior division | −2 | −38 | 36 | 6.57 | 15,072 |
| Left lateral occipital cortex, superior division | −48 | −70 | 30 | 4.59 | 5,568 |
| Left superior frontal gyrus | −28 | 24 | 54 | 4.48 | 5,312 |
| Left supramarginal gyrus, posterior division | −34 | −48 | 24 | 3.87 | 408 |
| Right lateral occipital cortex, superior division | 36 | −64 | 42 | 3.85 | 1,320 |
| Left precuneus cortex | −18 | −58 | 36 | 3.83 | 1,008 |
| Left lateral occipital cortex, superior division | −36 | −64 | 56 | 3.69 | 3,616 |
| Right precentral gyrus | 8 | −30 | 58 | 3.68 | 1,944 |
| Left cerebellum, lobules IV–V | −6 | −48 | −14 | 3.67 | 816 |
| Right middle frontal gyrus | 28 | 28 | 50 | 3.60 | 936 |
| Right lateral occipital cortex, superior division | 30 | −58 | 62 | 3.31 | 1,152 |
| Right precentral gyrus | 34 | −18 | 68 | 3.28 | 416 |
| Left postcentral gyrus | −28 | −38 | 64 | 3.20 | 1,040 |
| *Partner feedback, Loss > Gain (209 voxels; minimum surviving z = 2.71)* |  |  |  |  |  |
| Left superior frontal gyrus | 0 | 10 | 60 | 4.19 | 1,216 |
| Right superior temporal gyrus, posterior division | 48 | −24 | −4 | 3.28 | 456 |
| *Partner feedback, Gain > Loss (30,117 voxels; minimum surviving z = 2.70)* |  |  |  |  |  |
| Left accumbens | −8 | 8 | −6 | 7.69 | 102,240 |
| Right accumbens | 12 | 10 | −6 | 7.57 |  |
| Left cingulate gyrus, anterior division | −4 | 38 | −4 | 7.01 |  |
| Left putamen | −20 | 2 | −12 | 6.87 |  |
| Left cingulate gyrus, posterior division | −2 | −38 | 40 | 5.58 | 19,384 |
| Left superior frontal gyrus | −22 | 10 | 62 | 5.56 | 17,152 |
| Right cerebellum, Crus II | 48 | −64 | −42 | 5.16 | 26,352 |
| Left inferior temporal gyrus, temporooccipital part | −60 | −44 | −18 | 5.14 | 8,136 |
| Right occipital fusiform gyrus | 18 | −84 | −2 | 4.97 | 18,600 |
| Left lateral occipital cortex, superior division | −38 | −78 | 32 | 4.84 | 12,984 |
| Left inferior temporal gyrus, temporooccipital part | −46 | −46 | −36 | 4.76 | 7,352 |
| Right central opercular cortex | 62 | −12 | 10 | 4.38 | 4,096 |
| Right hippocampus | 30 | −14 | −22 | 4.37 | 1,760 |
| Right insular cortex | 44 | 0 | 2 | 4.16 | 440 |
| Left postcentral gyrus | −42 | −14 | 28 | 4.11 | 648 |
| Right middle temporal gyrus, posterior division | 64 | −8 | −22 | 4.08 | 1,040 |
| Right hippocampus | 28 | −38 | 0 | 3.90 | 912 |
| Left cerebellum, lobule IX | −12 | −48 | −38 | 3.88 | 456 |
| Right postcentral gyrus | 30 | −30 | 30 | 3.88 | 608 |
| Right frontal orbital cortex | 24 | 26 | −16 | 3.84 | 1,720 |
| Right postcentral gyrus | 48 | −30 | 58 | 3.84 | 1,248 |
| Right frontal pole | 16 | 68 | 4 | 3.79 | 704 |
| Right cerebellum, Crus II | 16 | −88 | −38 | 3.72 | 2,024 |
| Left cerebellum, Crus II | −16 | −90 | −34 | 3.63 | 592 |
| Left superior parietal lobule | −28 | −48 | 62 | 3.56 | 512 |
| Right frontal pole | 54 | 40 | 4 | 3.56 | 1,224 |
| Right lateral occipital cortex, superior division | 40 | −82 | 30 | 3.42 | 888 |
| Right superior parietal lobule | 36 | −46 | 68 | 3.36 | 616 |
| Right lateral occipital cortex, superior division | 14 | −74 | 46 | 3.29 | 496 |
| *Partner trust decision, Distrust > Trust (82,199 voxels; minimum surviving z = 2.31)* |  |  |  |  |  |
| Left inferior temporal gyrus, temporooccipital part | −48 | −60 | −16 | 7.83 | 355,520 |
| Left superior parietal lobule | −34 | −52 | 42 | 7.81 |  |
| Left lateral occipital cortex, superior division | −32 | −60 | 48 | 7.75 |  |
| Left superior parietal lobule | −46 | −40 | 52 | 7.74 |  |
| Left superior frontal gyrus | −4 | 22 | 50 | 7.80 | 299,680 |
| Left inferior frontal gyrus, pars opercularis | −36 | 12 | 26 | 7.77 |  |
| Left inferior frontal gyrus, pars opercularis | −46 | 10 | 28 | 7.53 |  |
| Left superior frontal gyrus | −2 | 28 | 54 | 7.12 |  |
| Left temporal pole | −44 | 2 | −36 | 4.12 | 1,120 |
| Right putamen | 34 | −10 | −10 | 3.81 | 648 |
| *Partner trust decision, Trust > Distrust (10,426 voxels; minimum surviving z = 2.31)* |  |  |  |  |  |
| Right hippocampus | 32 | −44 | 2 | 5.63 | 7,840 |
| Left hippocampus | −20 | −42 | 16 | 4.90 | 28,672 |
| Right parietal operculum cortex | 46 | −34 | 20 | 4.90 | 12,648 |
| Left paracingulate gyrus | −6 | 36 | −12 | 4.78 | 3,520 |
| Right postcentral gyrus | 22 | −40 | 62 | 4.75 | 13,368 |
| Left postcentral gyrus | −38 | −22 | 42 | 4.06 | 696 |
| Left central opercular cortex | −42 | −14 | 20 | 4.03 | 584 |
| Right cerebellum, lobule IX | 10 | −38 | −48 | 3.98 | 5,320 |
| Left hippocampus | −22 | −16 | −20 | 3.90 | 704 |
| Right hippocampus | 24 | −16 | −16 | 3.82 | 1,672 |
| Left lingual gyrus | −10 | −78 | −4 | 3.45 | 408 |
| Right cingulate gyrus, posterior division | 4 | −30 | 46 | 3.36 | 1,048 |
| Left precuneus cortex | −10 | −54 | 18 | 3.32 | 504 |
| Left postcentral gyrus | −22 | −32 | 58 | 2.92 | 600 |
| Left occipital pole | 0 | −100 | 12 | 2.80 | 1,080 |
| *Partner group difference, Human > System on Distrust > Trust (45,982 voxels; minimum surviving z = 2.56)* |  |  |  |  |  |
| Left inferior frontal gyrus, pars opercularis | −36 | 12 | 24 | 5.35 | 286,552 |
| Right cerebellum, lobule VIII | 30 | −62 | −46 | 5.18 |  |
| Right cerebellum, lobule VIII | 34 | −56 | −42 | 4.80 |  |
| Left lateral occipital cortex, superior division | −30 | −68 | 52 | 4.78 |  |
| Right angular gyrus | 44 | −46 | 48 | 4.73 | 10,176 |
| Right inferior frontal gyrus, pars opercularis | 34 | 14 | 24 | 4.66 | 6,000 |
| Left thalamus | −10 | −2 | 10 | 4.48 | 21,200 |
| Left superior frontal gyrus | −14 | 18 | 66 | 4.39 | 10,928 |
| Right frontal pole | 38 | 40 | 10 | 4.25 | 8,048 |
| Right supramarginal gyrus, posterior division | 58 | −40 | 34 | 4.06 | 1,336 |
| Right temporal fusiform cortex, anterior division | 42 | −6 | −30 | 4.02 | 1,152 |
| Left temporal pole | −46 | 20 | −38 | 3.99 | 952 |
| Left cerebellum, lobules IV–V | −16 | −32 | −26 | 3.86 | 1,136 |
| Right temporal pole | 54 | 20 | −22 | 3.79 | 1,120 |
| Right frontal pole | 34 | 40 | 40 | 3.73 | 3,032 |
| Right frontal orbital cortex | 26 | 30 | −14 | 3.62 | 872 |
| Left cingulate gyrus, anterior division | −6 | −6 | 34 | 3.61 | 1,024 |
| Right precuneus cortex | 14 | −50 | 26 | 3.55 | 848 |
| Right postcentral gyrus | 18 | −40 | 58 | 3.53 | 456 |
| Left amygdala | −10 | −14 | −12 | 3.50 | 1,056 |
| Left putamen | −34 | −22 | −6 | 3.48 | 528 |
| Left cingulate gyrus, posterior division | 0 | −34 | 32 | 3.42 | 672 |
| Left frontal pole | −10 | 52 | −16 | 3.41 | 1,016 |
| Right superior frontal gyrus | 16 | 30 | 52 | 3.31 | 1,024 |
| Left supplementary motor cortex | −16 | −10 | 46 | 3.30 | 440 |
| Right precentral gyrus | 38 | −4 | 56 | 3.18 | 504 |

*Note.* Second-level maps thresholded with voxel-wise false discovery rate correction (*q* < .05). Each heading gives the number of suprathreshold voxels and the smallest surviving z for that map. Clusters of at least 400 mm³ (50 voxels) are listed; for clusters larger than 50,000 mm³, up to three further local maxima at least 8 mm apart are listed, indented, beneath the cluster peak. Coordinates are in MNI152 space, and peak z is given as a positive value in the direction of each contrast. Labels are from the Harvard-Oxford atlas, with the AAL atlas used for cerebellum; where a peak lay outside labelled grey matter, the nearest label within 8 mm is given. Feedback and trust decision contrasts are computed across all participants (*N* = 44); the partner group difference is the Human > System contrast on Distrust > Trust.

**Table S2.** Decoding of Trust Versus Distrust Decisions Against Chance.

| ROI group | Sample | *n* | *M* (*SD*) | 95% CI | *t* | *df* | *q* | *d* |
|---|---|---|---|---|---|---|---|---|
| Whole task |  |  |  |  |  |  |  |  |
| Full network | Pooled | 42 | .643 (.135) | [.601, .685] | 6.90 | 41 | < .001 | 1.06 |
|  | Human | 20 | .698 (.126) | [.639, .758] | 7.01 | 19 | < .001 | 1.57 |
|  | System | 22 | .593 (.124) | [.538, .648] | 3.53 | 21 | .003 | 0.75 |
| Valuation | Pooled | 42 | .622 (.108) | [.589, .656] | 7.35 | 41 | < .001 | 1.13 |
|  | Human | 20 | .666 (.108) | [.615, .716] | 6.86 | 19 | < .001 | 1.53 |
|  | System | 22 | .583 (.093) | [.542, .624] | 4.17 | 21 | < .001 | 0.89 |
| Salience | Pooled | 42 | .633 (.118) | [.596, .670] | 7.30 | 41 | < .001 | 1.13 |
|  | Human | 20 | .682 (.120) | [.625, .738] | 6.75 | 19 | < .001 | 1.51 |
|  | System | 22 | .589 (.099) | [.545, .633] | 4.21 | 21 | < .001 | 0.90 |
| Social (rSTS) | Pooled | 42 | .599 (.102) | [.568, .631] | 6.33 | 41 | < .001 | 0.98 |
|  | Human | 20 | .636 (.108) | [.586, .686] | 5.65 | 19 | < .001 | 1.26 |
|  | System | 22 | .566 (.086) | [.528, .604] | 3.64 | 21 | .002 | 0.78 |
| Amygdala | Pooled | 42 | .552 (.070) | [.530, .573] | 4.79 | 41 | < .001 | 0.74 |
|  | Human | 20 | .584 (.080) | [.546, .621] | 4.68 | 19 | < .001 | 1.05 |
|  | System | 22 | .523 (.044) | [.503, .542] | 2.42 | 21 | .028 | 0.52 |
| Easy trials |  |  |  |  |  |  |  |  |
| Full network | Pooled | 40 | .802 (.163) | [.750, .854] | 11.73 | 39 | < .001 | 1.85 |
|  | Human | 19 | .818 (.166) | [.737, .898] | 8.32 | 18 | < .001 | 1.91 |
|  | System | 21 | .788 (.163) | [.714, .862] | 8.12 | 20 | < .001 | 1.77 |
| Valuation | Pooled | 40 | .796 (.180) | [.738, .853] | 10.38 | 39 | < .001 | 1.64 |
|  | Human | 19 | .810 (.193) | [.717, .903] | 6.99 | 18 | < .001 | 1.60 |
|  | System | 21 | .783 (.171) | [.705, .861] | 7.58 | 20 | < .001 | 1.65 |
| Salience | Pooled | 40 | .811 (.173) | [.756, .866] | 11.39 | 39 | < .001 | 1.80 |
|  | Human | 19 | .837 (.163) | [.759, .916] | 9.00 | 18 | < .001 | 2.06 |
|  | System | 21 | .787 (.181) | [.704, .869] | 7.26 | 20 | < .001 | 1.58 |
| Social (rSTS) | Pooled | 40 | .774 (.171) | [.719, .828] | 10.13 | 39 | < .001 | 1.60 |
|  | Human | 19 | .794 (.173) | [.711, .878] | 7.41 | 18 | < .001 | 1.70 |
|  | System | 21 | .755 (.171) | [.677, .833] | 6.84 | 20 | < .001 | 1.49 |
| Amygdala | Pooled | 40 | .758 (.165) | [.705, .810] | 9.86 | 39 | < .001 | 1.56 |
|  | Human | 19 | .793 (.156) | [.718, .868] | 8.20 | 18 | < .001 | 1.88 |
|  | System | 21 | .726 (.171) | [.648, .804] | 6.06 | 20 | < .001 | 1.32 |
| Hard trials |  |  |  |  |  |  |  |  |
| Full network | Pooled | 42 | .602 (.100) | [.571, .633] | 6.62 | 41 | < .001 | 1.02 |
|  | Human | 20 | .650 (.110) | [.599, .701] | 6.11 | 19 | < .001 | 1.37 |
|  | System | 22 | .558 (.066) | [.529, .587] | 4.14 | 21 | < .001 | 0.88 |
| Valuation | Pooled | 42 | .593 (.114) | [.558, .629] | 5.29 | 41 | < .001 | 0.82 |
|  | Human | 20 | .633 (.132) | [.572, .695] | 4.52 | 19 | < .001 | 1.01 |
|  | System | 22 | .556 (.082) | [.520, .593] | 3.23 | 21 | .006 | 0.69 |
| Salience | Pooled | 42 | .594 (.107) | [.560, .627] | 5.67 | 41 | < .001 | 0.87 |
|  | Human | 20 | .634 (.125) | [.575, .692] | 4.78 | 19 | < .001 | 1.07 |
|  | System | 22 | .557 (.072) | [.525, .589] | 3.70 | 21 | .002 | 0.79 |
| Social (rSTS) | Pooled | 42 | .572 (.100) | [.540, .603] | 4.64 | 41 | < .001 | 0.72 |
|  | Human | 20 | .595 (.111) | [.543, .647] | 3.83 | 19 | .002 | 0.86 |
|  | System | 22 | .550 (.086) | [.512, .589] | 2.74 | 21 | .015 | 0.58 |
| Amygdala | Pooled | 42 | .566 (.086) | [.539, .593] | 4.95 | 41 | < .001 | 0.76 |
|  | Human | 20 | .602 (.100) | [.555, .649] | 4.56 | 19 | < .001 | 1.02 |
|  | System | 22 | .533 (.056) | [.508, .558] | 2.77 | 21 | .014 | 0.59 |

*Note.* One-sample *t* tests against chance (.50). Pooled = both partner conditions combined; because group sizes differ, pooled means are weighted averages of the human and system means rather than an independent estimate. *q* = Benjamini–Hochberg false discovery rate, corrected across all 60 tests reported in Table 3 and Table S2. *d* = Cohen’s *d* for one sample. Decoding exceeded chance in all 45 cells.

**Table S3.** Within-Group Beta-Series Coupling for All 45 ROI Pairs, Hard Trust Trials.

|  | Human partner (*n* = 20) |  |  | System partner (*n* = 22) |  |  |
|---|---|---|---|---|---|---|
| ROI pair | *z* | *t* | *q* | *z* | *t* | *q* |
| aIns_L–aIns_R | 0.635 | 9.95 | < .001 | 0.631 | 12.87 | < .001 |
| SMA–aIns_L | 0.572 | 14.86 | < .001 | 0.543 | 8.18 | < .001 |
| SMA–aIns_R | 0.521 | 11.96 | < .001 | 0.440 | 7.04 | < .001 |
| aMCC–aIns_L | 0.472 | 9.78 | < .001 | 0.480 | 11.33 | < .001 |
| SMA–aMCC | 0.478 | 10.36 | < .001 | 0.405 | 7.01 | < .001 |
| rIFG–aIns_R | 0.425 | 6.06 | < .001 | 0.382 | 7.98 | < .001 |
| rIFG–aIns_L | 0.358 | 5.66 | < .001 | 0.441 | 10.34 | < .001 |
| rIFG–SMA | 0.385 | 9.13 | < .001 | 0.324 | 6.29 | < .001 |
| SMA–rSTS | 0.409 | 9.06 | < .001 | 0.292 | 4.80 | < .001 |
| aMCC–aIns_R | 0.399 | 8.35 | < .001 | 0.396 | 8.49 | < .001 |
| aIns_L–rSTS | 0.339 | 6.59 | < .001 | 0.356 | 8.93 | < .001 |
| aMCC–rSTS | 0.346 | 7.12 | < .001 | 0.339 | 7.53 | < .001 |
| vStr_L–vStr_R | 0.343 | 6.70 | < .001 | 0.366 | 9.92 | < .001 |
| aIns_R–rSTS | 0.301 | 6.81 | < .001 | 0.320 | 6.59 | < .001 |
| lOFC–rIFG | 0.311 | 5.94 | < .001 | 0.275 | 7.03 | < .001 |
| rIFG–aMCC | 0.313 | 6.13 | < .001 | 0.334 | 9.31 | < .001 |
| lOFC–SMA | 0.297 | 7.98 | < .001 | 0.298 | 5.54 | < .001 |
| rIFG–rSTS | 0.216 | 5.08 | < .001 | 0.314 | 7.43 | < .001 |
| lOFC–aMCC | 0.288 | 4.78 | < .001 | 0.245 | 6.61 | < .001 |
| lOFC–aIns_L | 0.235 | 5.84 | < .001 | 0.258 | 5.53 | < .001 |
| SMA–lAmyg | 0.237 | 6.70 | < .001 | 0.251 | 6.02 | < .001 |
| vStr_L–aMCC | 0.278 | 4.64 | < .001 | 0.230 | 5.79 | < .001 |
| aIns_L–lAmyg | 0.226 | 6.09 | < .001 | 0.244 | 6.07 | < .001 |
| aMCC–lAmyg | 0.232 | 5.60 | < .001 | 0.229 | 6.52 | < .001 |
| vStr_L–aIns_L | 0.227 | 6.14 | < .001 | 0.262 | 6.25 | < .001 |
| vStr_L–rSTS | 0.220 | 5.02 | < .001 | 0.239 | 7.33 | < .001 |
| lOFC–lAmyg | 0.231 | 5.87 | < .001 | 0.216 | 6.33 | < .001 |
| vStr_L–lAmyg | 0.264 | 8.77 | < .001 | 0.202 | 5.34 | < .001 |
| lOFC–rSTS | 0.202 | 4.35 | < .001 | 0.246 | 6.62 | < .001 |
| vStr_L–SMA | 0.238 | 5.66 | < .001 | 0.183 | 3.98 | < .001 |
| lOFC–vStr_R | 0.217 | 3.83 | .001 | 0.245 | 8.03 | < .001 |
| rSTS–lAmyg | 0.222 | 5.50 | < .001 | 0.241 | 4.87 | < .001 |
| vStr_R–aMCC | 0.203 | 3.70 | .002 | 0.241 | 5.95 | < .001 |
| rIFG–lAmyg | 0.212 | 8.15 | < .001 | 0.247 | 5.97 | < .001 |
| vStr_L–aIns_R | 0.220 | 5.26 | < .001 | 0.236 | 5.66 | < .001 |
| lOFC–vStr_L | 0.253 | 3.79 | .001 | 0.213 | 5.76 | < .001 |
| vStr_R–aIns_L | 0.173 | 4.14 | < .001 | 0.246 | 5.89 | < .001 |
| vStr_R–SMA | 0.175 | 4.85 | < .001 | 0.195 | 4.58 | < .001 |
| rIFG–vStr_R | 0.169 | 3.69 | .002 | 0.246 | 6.28 | < .001 |
| vStr_R–lAmyg | 0.172 | 5.78 | < .001 | 0.214 | 6.14 | < .001 |
| vStr_R–rSTS | 0.175 | 4.34 | < .001 | 0.209 | 5.60 | < .001 |
| rIFG–vStr_L | 0.215 | 5.01 | < .001 | 0.257 | 6.24 | < .001 |
| aIns_R–lAmyg | 0.188 | 4.93 | < .001 | 0.226 | 6.00 | < .001 |
| lOFC–aIns_R | 0.203 | 4.14 | < .001 | 0.175 | 3.97 | < .001 |
| vStr_R–aIns_R | 0.186 | 4.35 | < .001 | 0.191 | 4.16 | < .001 |

*Note.* Values are mean Fisher *z*-transformed correlations tested against zero with one-sample *t* tests. Human *n* = 20 (*df* = 19), system *n* = 22 (*df* = 21). *q* values are false discovery rate corrected (Benjamini–Hochberg) across the 45 pairs within each group. Rows are ordered by descending mean coupling across both decision types and both partner groups, so Tables S3 and S4 share one row order.

**Table S4.** Within-Group Beta-Series Coupling for All 45 ROI Pairs, Hard Distrust Trials.

|  | Human partner (*n* = 20) |  |  | System partner (*n* = 22) |  |  |
|---|---|---|---|---|---|---|
| ROI pair | *z* | *t* | *q* | *z* | *t* | *q* |
| aIns_L–aIns_R | 0.673 | 7.52 | < .001 | 0.621 | 12.45 | < .001 |
| SMA–aIns_L | 0.663 | 9.80 | < .001 | 0.544 | 9.10 | < .001 |
| SMA–aIns_R | 0.582 | 8.24 | < .001 | 0.445 | 6.81 | < .001 |
| aMCC–aIns_L | 0.405 | 5.67 | < .001 | 0.404 | 7.45 | < .001 |
| SMA–aMCC | 0.461 | 5.45 | < .001 | 0.401 | 7.19 | < .001 |
| rIFG–aIns_R | 0.471 | 4.72 | < .001 | 0.379 | 8.77 | < .001 |
| rIFG–aIns_L | 0.389 | 4.56 | < .001 | 0.435 | 10.65 | < .001 |
| rIFG–SMA | 0.499 | 6.46 | < .001 | 0.371 | 7.30 | < .001 |
| SMA–rSTS | 0.454 | 7.65 | < .001 | 0.321 | 5.86 | < .001 |
| aMCC–aIns_R | 0.318 | 4.41 | < .001 | 0.323 | 7.43 | < .001 |
| aIns_L–rSTS | 0.358 | 5.37 | < .001 | 0.345 | 9.00 | < .001 |
| aMCC–rSTS | 0.368 | 4.99 | < .001 | 0.281 | 7.13 | < .001 |
| vStr_L–vStr_R | 0.301 | 3.75 | .002 | 0.321 | 8.15 | < .001 |
| aIns_R–rSTS | 0.378 | 5.86 | < .001 | 0.310 | 7.68 | < .001 |
| lOFC–rIFG | 0.436 | 5.72 | < .001 | 0.223 | 7.14 | < .001 |
| rIFG–aMCC | 0.311 | 5.01 | < .001 | 0.284 | 6.71 | < .001 |
| lOFC–SMA | 0.396 | 5.31 | < .001 | 0.234 | 6.37 | < .001 |
| rIFG–rSTS | 0.278 | 4.44 | < .001 | 0.282 | 9.94 | < .001 |
| lOFC–aMCC | 0.328 | 3.87 | .002 | 0.190 | 6.35 | < .001 |
| lOFC–aIns_L | 0.317 | 3.92 | .002 | 0.223 | 5.24 | < .001 |
| SMA–lAmyg | 0.255 | 3.11 | .007 | 0.281 | 5.57 | < .001 |
| vStr_L–aMCC | 0.260 | 4.09 | .001 | 0.224 | 6.95 | < .001 |
| aIns_L–lAmyg | 0.241 | 3.90 | .002 | 0.238 | 6.82 | < .001 |
| aMCC–lAmyg | 0.258 | 3.38 | .004 | 0.224 | 4.63 | < .001 |
| vStr_L–aIns_L | 0.237 | 3.43 | .004 | 0.207 | 6.29 | < .001 |
| vStr_L–rSTS | 0.266 | 6.03 | < .001 | 0.202 | 5.24 | < .001 |
| lOFC–lAmyg | 0.326 | 5.10 | < .001 | 0.152 | 3.31 | .003 |
| vStr_L–lAmyg | 0.234 | 3.14 | .006 | 0.210 | 6.74 | < .001 |
| lOFC–rSTS | 0.246 | 4.69 | < .001 | 0.212 | 7.00 | < .001 |
| vStr_L–SMA | 0.279 | 3.84 | .002 | 0.203 | 6.73 | < .001 |
| lOFC–vStr_R | 0.263 | 3.79 | .002 | 0.172 | 6.41 | < .001 |
| rSTS–lAmyg | 0.254 | 3.55 | .003 | 0.177 | 4.79 | < .001 |
| vStr_R–aMCC | 0.188 | 2.62 | .017 | 0.230 | 6.27 | < .001 |
| rIFG–lAmyg | 0.176 | 3.07 | .007 | 0.217 | 6.19 | < .001 |
| vStr_L–aIns_R | 0.207 | 2.89 | .010 | 0.187 | 3.98 | < .001 |
| lOFC–vStr_L | 0.189 | 2.24 | .037 | 0.193 | 7.59 | < .001 |
| vStr_R–aIns_L | 0.228 | 3.80 | .002 | 0.196 | 5.53 | < .001 |
| vStr_R–SMA | 0.275 | 5.77 | < .001 | 0.198 | 5.39 | < .001 |
| rIFG–vStr_R | 0.185 | 3.66 | .002 | 0.228 | 8.47 | < .001 |
| vStr_R–lAmyg | 0.231 | 3.45 | .004 | 0.203 | 5.50 | < .001 |
| vStr_R–rSTS | 0.207 | 3.17 | .006 | 0.225 | 4.96 | < .001 |
| rIFG–vStr_L | 0.161 | 2.70 | .015 | 0.177 | 5.09 | < .001 |
| aIns_R–lAmyg | 0.171 | 3.92 | .002 | 0.204 | 5.69 | < .001 |
| lOFC–aIns_R | 0.213 | 2.84 | .011 | 0.168 | 4.41 | < .001 |
| vStr_R–aIns_R | 0.181 | 3.13 | .006 | 0.164 | 4.81 | < .001 |

*Note.* Values are mean Fisher *z*-transformed correlations tested against zero with one-sample *t* tests. Human *n* = 20 (*df* = 19), system *n* = 22 (*df* = 21). *q* values are false discovery rate corrected (Benjamini–Hochberg) across the 45 pairs within each group. Rows are ordered by descending mean coupling across both decision types and both partner groups, so Tables S3 and S4 share one row order.

**Table S5.** Between-Group Contrast (Human − System) for All 45 ROI Pairs, Hard Trust Trials.

| ROI pair | Human *z* | System *z* | Δ*z* | 95% CI | *d* | *t* | *df* | *p* | *q* |
|---|---|---|---|---|---|---|---|---|---|
| rIFG–rSTS | 0.216 | 0.314 | −0.097 | [−0.218, +0.024] | −0.50 | −1.62 | 39.9 | .112 | .944 |
| SMA–rSTS | 0.409 | 0.292 | +0.118 | [−0.036, +0.271] | 0.47 | 1.55 | 37.9 | .129 | .944 |
| vStr_L–lAmyg | 0.264 | 0.202 | +0.062 | [−0.036, +0.160] | 0.39 | 1.28 | 38.8 | .208 | .944 |
| rIFG–vStr_R | 0.169 | 0.246 | −0.076 | [−0.198, +0.045] | −0.39 | −1.27 | 38.4 | .212 | .944 |
| vStr_R–aIns_L | 0.173 | 0.246 | −0.074 | [−0.193, +0.046] | −0.38 | −1.25 | 39.9 | .219 | .944 |
| rIFG–aIns_L | 0.358 | 0.441 | −0.082 | [−0.237, +0.073] | −0.34 | −1.08 | 33.8 | .289 | .944 |
| SMA–aIns_R | 0.521 | 0.440 | +0.082 | [−0.073, +0.236] | 0.33 | 1.07 | 36.8 | .290 | .944 |
| SMA–aMCC | 0.478 | 0.405 | +0.073 | [−0.077, +0.222] | 0.30 | 0.98 | 38.9 | .332 | .944 |
| vStr_R–lAmyg | 0.172 | 0.214 | −0.042 | [−0.135, +0.051] | −0.28 | −0.92 | 39.5 | .364 | .944 |
| rIFG–SMA | 0.385 | 0.324 | +0.060 | [−0.074, +0.195] | 0.28 | 0.91 | 39.1 | .369 | .944 |
| vStr_L–SMA | 0.238 | 0.183 | +0.054 | [−0.072, +0.180] | 0.27 | 0.87 | 39.9 | .390 | .944 |
| lOFC–rSTS | 0.202 | 0.246 | −0.044 | [−0.164, +0.077] | −0.23 | −0.74 | 37.3 | .466 | .944 |
| rIFG–lAmyg | 0.212 | 0.247 | −0.035 | [−0.135, +0.064] | −0.22 | −0.72 | 34.8 | .474 | .944 |
| rIFG–vStr_L | 0.215 | 0.257 | −0.042 | [−0.162, +0.078] | −0.22 | −0.70 | 39.7 | .485 | .944 |
| aIns_R–lAmyg | 0.188 | 0.226 | −0.038 | [−0.146, +0.071] | −0.22 | −0.70 | 39.8 | .486 | .944 |
| vStr_L–aMCC | 0.278 | 0.231 | +0.047 | [−0.099, +0.194] | 0.21 | 0.66 | 33.6 | .514 | .944 |
| vStr_L–aIns_L | 0.227 | 0.262 | −0.035 | [−0.148, +0.078] | −0.19 | −0.63 | 39.8 | .531 | .944 |
| vStr_R–rSTS | 0.175 | 0.209 | −0.035 | [−0.146, +0.077] | −0.19 | −0.63 | 39.4 | .533 | .944 |
| lOFC–aMCC | 0.287 | 0.245 | +0.043 | [−0.101, +0.187] | 0.19 | 0.61 | 32.0 | .547 | .944 |
| vStr_R–aMCC | 0.203 | 0.241 | −0.038 | [−0.177, +0.100] | −0.18 | −0.56 | 35.8 | .579 | .944 |
| lOFC–rIFG | 0.311 | 0.275 | +0.036 | [−0.097, +0.168] | 0.17 | 0.55 | 36.0 | .587 | .944 |
| lOFC–vStr_L | 0.253 | 0.213 | +0.040 | [−0.116, +0.196] | 0.17 | 0.53 | 29.9 | .601 | .944 |
| rIFG–aIns_R | 0.425 | 0.382 | +0.043 | [−0.129, +0.216] | 0.16 | 0.51 | 34.1 | .612 | .944 |
| lOFC–aIns_R | 0.203 | 0.175 | +0.028 | [−0.105, +0.161] | 0.13 | 0.43 | 39.0 | .671 | .944 |
| lOFC–vStr_R | 0.217 | 0.245 | −0.028 | [−0.159, +0.104] | −0.14 | −0.43 | 29.3 | .672 | .944 |
| SMA–aIns_L | 0.572 | 0.543 | +0.029 | [−0.127, +0.185] | 0.11 | 0.38 | 33.3 | .706 | .944 |
| vStr_L–vStr_R | 0.343 | 0.366 | −0.024 | [−0.152, +0.104] | −0.12 | −0.38 | 35.3 | .710 | .944 |
| vStr_R–SMA | 0.175 | 0.195 | −0.021 | [−0.134, +0.092] | −0.11 | −0.37 | 39.4 | .713 | .944 |
| lOFC–aIns_L | 0.235 | 0.258 | −0.022 | [−0.147, +0.102] | −0.11 | −0.36 | 39.6 | .719 | .944 |
| vStr_L–rSTS | 0.220 | 0.239 | −0.019 | [−0.130, +0.091] | −0.11 | −0.35 | 35.9 | .726 | .944 |
| rIFG–aMCC | 0.313 | 0.334 | −0.021 | [−0.148, +0.105] | −0.11 | −0.34 | 34.8 | .732 | .944 |
| aIns_L–lAmyg | 0.226 | 0.244 | −0.018 | [−0.129, +0.093] | −0.10 | −0.33 | 40.0 | .743 | .944 |
| rSTS–lAmyg | 0.222 | 0.241 | −0.019 | [−0.149, +0.110] | −0.09 | −0.31 | 39.1 | .761 | .944 |
| aIns_R–rSTS | 0.301 | 0.320 | −0.019 | [−0.152, +0.114] | −0.09 | −0.29 | 39.9 | .773 | .944 |
| lOFC–lAmyg | 0.231 | 0.216 | +0.015 | [−0.091, +0.120] | 0.09 | 0.29 | 38.6 | .777 | .944 |
| vStr_L–aIns_R | 0.220 | 0.236 | −0.016 | [−0.135, +0.103] | −0.08 | −0.27 | 39.9 | .787 | .944 |
| SMA–lAmyg | 0.237 | 0.251 | −0.014 | [−0.125, +0.096] | −0.08 | −0.26 | 39.5 | .794 | .944 |
| aIns_L–rSTS | 0.339 | 0.356 | −0.017 | [−0.149, +0.115] | −0.08 | −0.26 | 36.7 | .797 | .944 |
| aMCC–aIns_L | 0.472 | 0.480 | −0.008 | [−0.138, +0.122] | −0.04 | −0.13 | 38.8 | .898 | .987 |
| aMCC–rSTS | 0.346 | 0.339 | +0.007 | [−0.127, +0.141] | 0.03 | 0.11 | 39.4 | .916 | .987 |
| vStr_R–aIns_R | 0.186 | 0.191 | −0.005 | [−0.132, +0.122] | −0.03 | −0.09 | 40.0 | .932 | .987 |
| aIns_L–aIns_R | 0.635 | 0.631 | +0.004 | [−0.159, +0.167] | 0.02 | 0.05 | 36.5 | .958 | .987 |
| aMCC–lAmyg | 0.232 | 0.229 | +0.002 | [−0.108, +0.112] | 0.01 | 0.04 | 38.3 | .967 | .987 |
| aMCC–aIns_R | 0.399 | 0.396 | +0.003 | [−0.133, +0.138] | 0.01 | 0.04 | 39.8 | .970 | .987 |
| lOFC–SMA | 0.297 | 0.298 | −0.001 | [−0.134, +0.131] | −0.00 | −0.02 | 36.6 | .987 | .987 |

*Note.* Δ*z* is the difference in mean Fisher *z*-transformed coupling (human minus system); positive values indicate stronger coupling in the human group. Human *n* = 20, system *n* = 22. Welch *t* tests were used because group sizes and variances were unequal; d is Cohen's *d* computed with the pooled standard deviation. *q* values are false discovery rate corrected (Benjamini–Hochberg) across the 45 pairs. Rows are ordered by ascending uncorrected p. No pair reached significance either uncorrected or after correction.

**Table S6.** Between-Group Contrast (Human − System) for All 45 ROI Pairs, Hard Distrust Trials.

| ROI pair | Human *z* | System *z* | Δ*z* | 95% CI | *d* | *t* | *df* | *p* | *q* |
|---|---|---|---|---|---|---|---|---|---|
| lOFC–rIFG* | 0.436 | 0.223 | +0.213 | [+0.043, +0.383] | 0.83 | 2.58 | 25.3 | .016 | .714 |
| lOFC–lAmyg* | 0.326 | 0.152 | +0.174 | [+0.015, +0.334] | 0.69 | 2.21 | 35.1 | .033 | .752 |
| lOFC–SMA | 0.396 | 0.234 | +0.162 | [−0.008, +0.333] | 0.62 | 1.95 | 27.8 | .061 | .917 |
| SMA–rSTS | 0.454 | 0.321 | +0.133 | [−0.030, +0.297] | 0.51 | 1.65 | 39.3 | .107 | .956 |
| lOFC–aMCC | 0.328 | 0.190 | +0.138 | [−0.048, +0.324] | 0.49 | 1.54 | 23.7 | .137 | .956 |
| SMA–aIns_R | 0.582 | 0.445 | +0.137 | [−0.058, +0.332] | 0.44 | 1.42 | 39.4 | .162 | .956 |
| rIFG–SMA | 0.499 | 0.371 | +0.128 | [−0.060, +0.316] | 0.43 | 1.38 | 33.3 | .176 | .956 |
| SMA–aIns_L | 0.663 | 0.544 | +0.119 | [−0.064, +0.302] | 0.41 | 1.32 | 38.8 | .195 | .956 |
| vStr_R–SMA | 0.275 | 0.198 | +0.077 | [−0.045, +0.199] | 0.40 | 1.28 | 36.6 | .208 | .956 |
| lOFC–vStr_R | 0.263 | 0.172 | +0.091 | [−0.062, +0.244] | 0.39 | 1.23 | 24.6 | .232 | .956 |
| vStr_L–rSTS | 0.267 | 0.202 | +0.065 | [−0.054, +0.183] | 0.34 | 1.11 | 38.6 | .275 | .956 |
| aMCC–rSTS | 0.368 | 0.281 | +0.087 | [−0.084, +0.258] | 0.33 | 1.04 | 29.2 | .307 | .956 |
| lOFC–aIns_L | 0.317 | 0.223 | +0.094 | [−0.093, +0.281] | 0.33 | 1.02 | 29.0 | .314 | .956 |
| vStr_L–SMA | 0.279 | 0.203 | +0.076 | [−0.086, +0.238] | 0.31 | 0.96 | 25.4 | .344 | .956 |
| rSTS–lAmyg | 0.254 | 0.177 | +0.077 | [−0.088, +0.243] | 0.30 | 0.96 | 28.6 | .345 | .956 |
| aIns_R–rSTS | 0.378 | 0.310 | +0.068 | [−0.087, +0.223] | 0.28 | 0.90 | 32.3 | .377 | .956 |
| rIFG–aIns_R | 0.471 | 0.379 | +0.092 | [−0.131, +0.316] | 0.27 | 0.85 | 26.0 | .403 | .956 |
| rIFG–vStr_R | 0.185 | 0.228 | −0.043 | [−0.161, +0.073] | −0.24 | −0.76 | 29.2 | .453 | .956 |
| rIFG–lAmyg | 0.176 | 0.217 | −0.041 | [−0.178, +0.096] | −0.19 | −0.61 | 31.9 | .544 | .956 |
| aIns_R–lAmyg | 0.171 | 0.204 | −0.034 | [−0.148, +0.081] | −0.18 | −0.59 | 37.8 | .556 | .956 |
| SMA–aMCC | 0.461 | 0.401 | +0.060 | [−0.146, +0.266] | 0.18 | 0.59 | 33.4 | .561 | .956 |
| lOFC–rSTS | 0.246 | 0.212 | +0.034 | [−0.089, +0.158] | 0.18 | 0.56 | 30.7 | .577 | .956 |
| lOFC–aIns_R | 0.213 | 0.168 | +0.045 | [−0.127, +0.217] | 0.17 | 0.54 | 28.4 | .596 | .956 |
| vStr_R–aMCC | 0.188 | 0.230 | −0.042 | [−0.206, +0.123] | −0.16 | −0.52 | 28.5 | .608 | .956 |
| aIns_L–aIns_R | 0.673 | 0.621 | +0.053 | [−0.157, +0.262] | 0.16 | 0.51 | 30.0 | .612 | .956 |
| vStr_L–aMCC | 0.260 | 0.224 | +0.035 | [−0.110, +0.181] | 0.16 | 0.50 | 28.4 | .622 | .956 |
| rIFG–aIns_L | 0.389 | 0.435 | −0.046 | [−0.240, +0.148] | −0.16 | −0.49 | 27.4 | .629 | .956 |
| vStr_R–aIns_L | 0.228 | 0.196 | +0.032 | [−0.110, +0.174] | 0.14 | 0.46 | 31.1 | .650 | .956 |
| vStr_L–aIns_L | 0.237 | 0.207 | +0.030 | [−0.127, +0.187] | 0.13 | 0.40 | 27.3 | .695 | .956 |
| aMCC–lAmyg | 0.258 | 0.224 | +0.035 | [−0.149, +0.218] | 0.12 | 0.38 | 32.5 | .705 | .956 |
| rIFG–aMCC | 0.311 | 0.284 | +0.027 | [−0.126, +0.180] | 0.11 | 0.36 | 34.1 | .719 | .956 |
| vStr_R–lAmyg | 0.231 | 0.203 | +0.027 | [−0.129, +0.183] | 0.11 | 0.36 | 29.9 | .724 | .956 |
| vStr_L–lAmyg | 0.234 | 0.210 | +0.023 | [−0.143, +0.189] | 0.09 | 0.29 | 25.6 | .776 | .956 |
| SMA–lAmyg | 0.255 | 0.281 | −0.026 | [−0.222, +0.170] | −0.09 | −0.27 | 31.9 | .788 | .956 |
| vStr_R–aIns_R | 0.181 | 0.164 | +0.017 | [−0.119, +0.154] | 0.08 | 0.26 | 31.1 | .797 | .956 |
| vStr_L–aIns_R | 0.207 | 0.187 | +0.020 | [−0.154, +0.195] | 0.07 | 0.24 | 33.2 | .813 | .956 |
| vStr_R–rSTS | 0.207 | 0.225 | −0.018 | [−0.180, +0.144] | −0.07 | −0.23 | 34.5 | .823 | .956 |
| vStr_L–vStr_R | 0.301 | 0.321 | −0.020 | [−0.203, +0.163] | −0.07 | −0.22 | 27.8 | .824 | .956 |
| rIFG–vStr_L | 0.162 | 0.177 | −0.015 | [−0.156, +0.126] | −0.07 | −0.22 | 30.8 | .828 | .956 |
| aIns_L–rSTS | 0.358 | 0.345 | +0.013 | [−0.144, +0.169] | 0.05 | 0.16 | 30.7 | .872 | .981 |
| aMCC–aIns_R | 0.318 | 0.323 | −0.005 | [−0.177, +0.166] | −0.02 | −0.06 | 31.6 | .950 | .990 |
| rIFG–rSTS | 0.278 | 0.282 | −0.004 | [−0.145, +0.137] | −0.02 | −0.06 | 26.6 | .954 | .990 |
| lOFC–vStr_L | 0.189 | 0.193 | −0.004 | [−0.186, +0.179] | −0.01 | −0.04 | 22.4 | .967 | .990 |
| aIns_L–lAmyg | 0.241 | 0.238 | +0.003 | [−0.142, +0.148] | 0.01 | 0.04 | 30.3 | .969 | .990 |
| aMCC–aIns_L | 0.405 | 0.404 | +0.001 | [−0.181, +0.183] | 0.00 | 0.01 | 36.3 | .990 | .990 |

*Note.* Δ*z* is the difference in mean Fisher *z*-transformed coupling (human minus system); positive values indicate stronger coupling in the human group. Human *n* = 20, system *n* = 22. Welch *t* tests were used because group sizes and variances were unequal; d is Cohen’s *d* computed with the pooled standard deviation. *q* values are false discovery rate corrected (Benjamini–Hochberg) across the 45 pairs. Rows are ordered by ascending uncorrected p. Asterisk indicates *p* < .05 uncorrected; no pair survived correction.

**Table S7.**
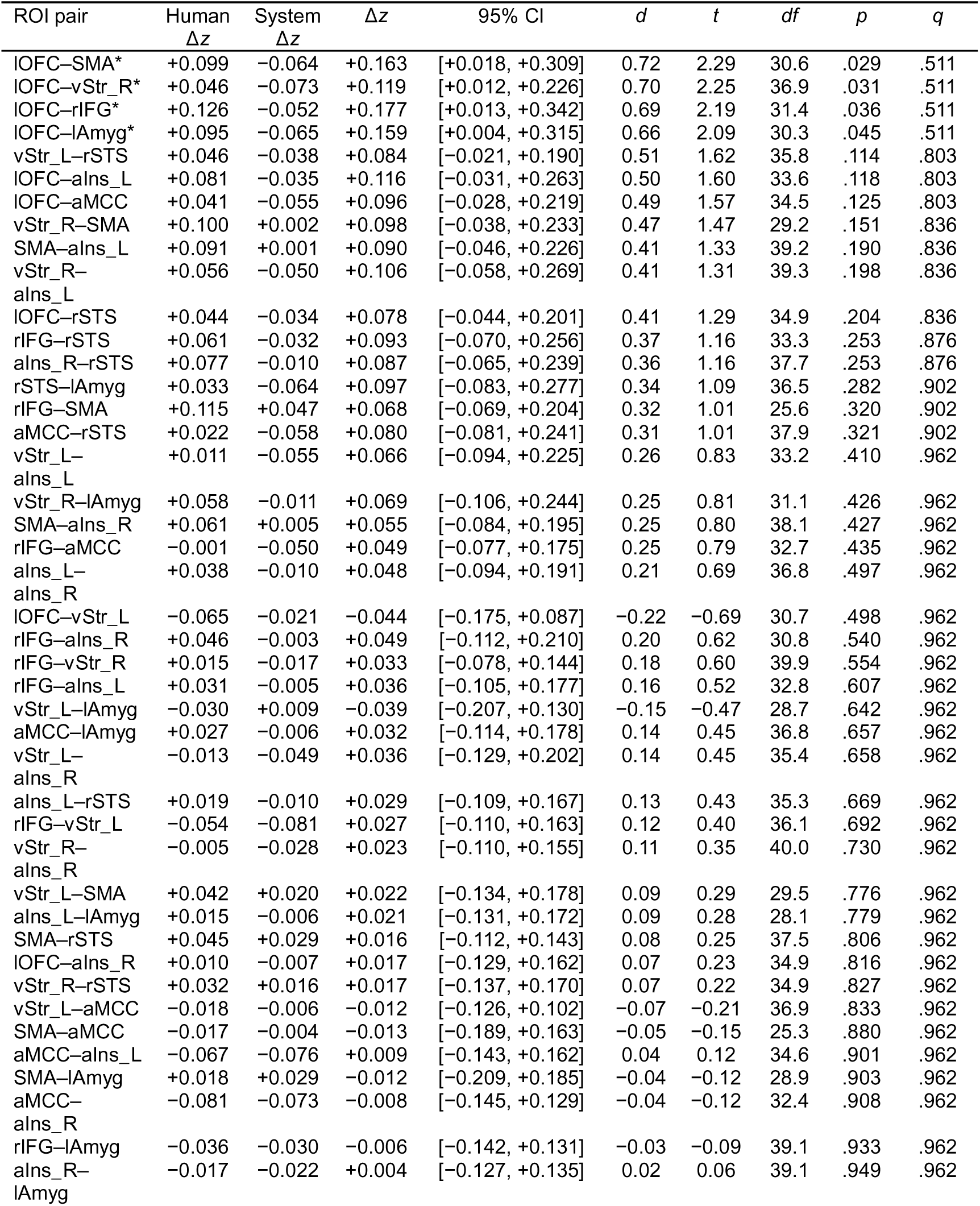
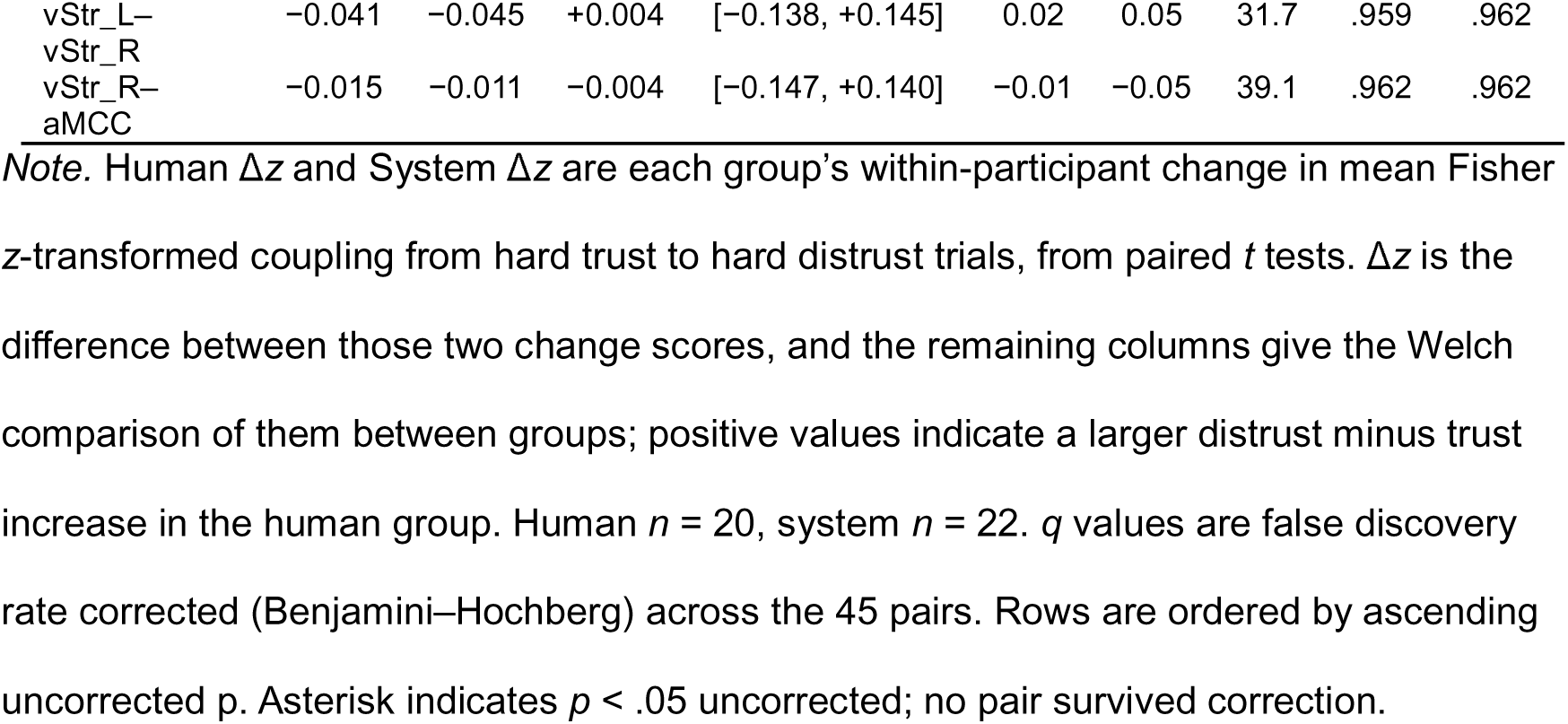
Within-Participant Distrust Minus Trust Coupling and the Partner Group Comparison, Hard Trials.

## References

Abdulrahman, H., & Henson, R. N. (2016). Effect of trial-to-trial variability on optimal event-related fMRI design: Implications for beta-series correlation and multi-voxel pattern analysis. NeuroImage, 125, 756–766. 10.1016/j.neuroimage.2015.11.009

Aben, B., Buc Calderon, C., Van den Bussche, E., & Verguts, T. (2020). Cognitive effort modulates connectivity between dorsal anterior cingulate cortex and task-relevant cortical areas. Journal of Neuroscience, 40(19), 3838–3848. 10.1523/JNEUROSCI.2948-19.2020

Abraham, A., Pedregosa, F., Eickenberg, M., Gervais, P., Mueller, A., Kossaifi, J., Gramfort, A., Thirion, B., & Varoquaux, G. (2014). Machine learning for neuroimaging with scikit-learn. Frontiers in Neuroinformatics, 8, 14. 10.3389/fninf.2014.00014

Aimone, J. A., Houser, D., & Weber, B. (2014). Neural signatures of betrayal aversion: An fMRI study of trust. Proceedings of the Royal Society B: Biological Sciences, 281(1782), 20132127. 10.1098/rspb.2013.2127

Bartra, O., McGuire, J. T., & Kable, J. W. (2013). The valuation system: A coordinate-based meta-analysis of BOLD fMRI experiments examining neural correlates of subjective value. NeuroImage, 76, 412–427. 10.1016/j.neuroimage.2013.02.063

Behrens, T. E. J., Hunt, L. T., Woolrich, M. W., & Rushworth, M. F. S. (2008). Associative learning of social value. Nature, 456(7219), 245–249. 10.1038/nature07538

Behzadi, Y., Restom, K., Liau, J., & Liu, T. T. (2007). A component based noise correction method (CompCor) for BOLD and perfusion based fMRI. NeuroImage, 37(1), 90–101. 10.1016/j.neuroimage.2007.04.042

Bellucci, G., Chernyak, S. V., Goodyear, K., Eickhoff, S. B., & Krueger, F. (2017). Neural signatures of trust in reciprocity: A coordinate-based meta-analysis. Human Brain Mapping, 38(3), 1233–1248. 10.1002/hbm.23451

Bellucci, G., Feng, C., Camilleri, J., Eickhoff, S. B., & Krueger, F. (2018). The role of the anterior insula in social norm compliance and enforcement: Evidence from coordinate-based and functional connectivity meta-analyses. Neuroscience & Biobehavioral Reviews, 92, 378–389. 10.1016/j.neubiorev.2018.06.024

Benjamini, Y., & Hochberg, Y. (1995). Controlling the false discovery rate: A practical and powerful approach to multiple testing. Journal of the Royal Statistical Society: Series B, 57(1), 289–300. 10.1111/j.2517-6161.1995.tb02031.x

Chaminade, T., Rosset, D., Da Fonseca, D., Nazarian, B., Lutcher, E., Cheng, G., & Deruelle, C. (2012). How do we think machines think? An fMRI study of alleged competition with an artificial intelligence. Frontiers in Human Neuroscience, 6, 103. 10.3389/fnhum.2012.00103

Chang, S. W. C., Fagan, N. A., Toda, K., Utevsky, A. V., Pearson, J. M., & Platt, M. L. (2015). Neural mechanisms of social decision-making in the primate amygdala. Proceedings of the National Academy of Sciences, 112(52), 16012–16017. 10.1073/pnas.1514761112

Chen, H., Chan, I. Y. S., Dong, Z., & Samuel, T.-A. (2025). Unraveling trust in collaborative human-machine intelligence from neurophysiological perspective: A review of EEG and fNIRS features. Advanced Engineering Informatics, 67, 103555. 10.1016/j.aei.2025.103555

Cox, R. W., & Hyde, J. S. (1997). Software tools for analysis and visualization of fMRI data. NMR in Biomedicine, 10(4–5), 171–178. 10.1002/(SICI)1099-1492(199706/08)10:4/5<171::AID-NBM453>3.0.CO;2-L

Craig, A. D. (2009). How do you feel - now? The anterior insula and human awareness. Nature Reviews Neuroscience, 10(1), 59–70. 10.1038/nrn2555

de Visser, E. J., Beatty, P. J., Estepp, J. R., Kohn, S., Abubshait, A., Fedota, J. R., & McDonald, C. G. (2018). Learning from the slips of others: Neural correlates of trust in automated agents. Frontiers in Human Neuroscience, 12, 309. 10.3389/fnhum.2018.00309

de Visser, E. J., Monfort, S. S., McKendrick, R., Smith, M. A. B., McKnight, P. E., Krueger, F., & Parasuraman, R. (2016). Almost human: Anthropomorphism increases trust resilience in cognitive agents. Journal of Experimental Psychology: Applied, 22(3), 331–349. 10.1037/xap0000092

Dietvorst, B. J., Simmons, J. P., & Massey, C. (2015). Algorithm aversion: People erroneously avoid algorithms after seeing them err. Journal of Experimental Psychology: General, 144(1), 114–126. 10.1037/xge0000033

Drnec, K., Marathe, A. R., Lukos, J. R., & Metcalfe, J. S. (2016). From trust in automation to decision neuroscience: Applying cognitive neuroscience methods to understand and improve interaction decisions involved in human automation interaction. Frontiers in Human Neuroscience, 10, 290. 10.3389/fnhum.2016.00290

Dzindolet, M. T., Peterson, S. A., Pomranky, R. A., Pierce, L. G., & Beck, H. P. (2003). The role of trust in automation reliance. International Journal of Human-Computer Studies, 58(6), 697–718. 10.1016/S1071-5819(03)00038-7

Esteban, O., Markiewicz, C. J., Blair, R. W., Moodie, C. A., Isik, A. I., Erramuzpe, A., Kent, J. D., Goncalves, M., DuPre, E., Snyder, M., Oya, H., Ghosh, S. S., Wright, J., Durnez, J., Poldrack, R. A., & Gorgolewski, K. J. (2019). fMRIPrep: A robust preprocessing pipeline for functional MRI. Nature Methods, 16(1), 111–116. 10.1038/s41592-018-0235-4

Gangopadhyay, P., Chawla, M., Dal Monte, O., & Chang, S. W. C. (2021). Prefrontal-amygdala circuits in social decision-making. Nature Neuroscience, 24(1), 5–18. 10.1038/s41593-020-00738-9

Glover, G. H. (1999). Deconvolution of impulse response in event-related BOLD fMRI. NeuroImage, 9(4), 416–429. 10.1006/nimg.1998.0419

Goodyear, K., Parasuraman, R., Chernyak, S., de Visser, E., Madhavan, P., Deshpande, G., & Krueger, F. (2017). An fMRI and effective connectivity study investigating miss errors during advice utilization from human and machine agents. Social Neuroscience, 12(5), 570–581. 10.1080/17470919.2016.1205131

Goodyear, K., Parasuraman, R., Chernyak, S., Madhavan, P., Deshpande, G., & Krueger, F. (2016). Advice taking from humans and machines: An fMRI and effective connectivity study. Frontiers in Human Neuroscience, 10, 542. 10.3389/fnhum.2016.00542

Greve, D. N., & Fischl, B. (2009). Accurate and robust brain image alignment using boundary-based registration. NeuroImage, 48(1), 63–72. 10.1016/j.neuroimage.2009.06.060

Grinband, J., Hirsch, J., & Ferrera, V. P. (2006). A neural representation of categorization uncertainty in the human brain. Neuron, 49(5), 757–763. 10.1016/j.neuron.2006.01.032

Hancock, P. A., Kessler, T. T., Kaplan, A. D., Brill, J. C., & Szalma, J. L. (2021). Evolving trust in robots: Specification through sequential and comparative meta-analyses. Human Factors, 63(7), 1196–1229. 10.1177/0018720820922080

Hoff, K. A., & Bashir, M. (2015). Trust in automation: Integrating empirical evidence on factors that influence trust. Human Factors, 57(3), 407–434. 10.1177/0018720814547570

Hogenhuis, A., & Hortensius, R. (2022). Domain-specific and domain-general neural network engagement during human-robot interactions. European Journal of Neuroscience, 56(10), 5902–5916. 10.1111/ejn.15823

Hopko, S. K., & Mehta, R. K. (2021). Neural correlates of trust in automation: Considerations and generalizability between technology domains. Frontiers in Neuroergonomics, 2, 731327. 10.3389/fnrgo.2021.731327

Hopko, S. K., Zhang, Y., Yadav, A., Pagilla, P. R., & Mehta, R. K. (2024). Brain-behavior relationships of trust in shared space human-robot collaboration. ACM Transactions on Human-Robot Interaction, 13(1), 1–23. 10.1145/3632149

Hortensius, R., & Cross, E. S. (2018). From automata to animate beings: The scope and limits of attributing socialness to artificial agents. Annals of the New York Academy of Sciences, 1426(1), 93–110. 10.1111/nyas.13727

Isik, L., Koldewyn, K., Beeler, D., & Kanwisher, N. (2017). Perceiving social interactions in the posterior superior temporal sulcus. Proceedings of the National Academy of Sciences, 114(43), E9145–E9152. 10.1073/pnas.1714471114

Jenkinson, M., & Smith, S. (2001). A global optimisation method for robust affine registration of brain images. Medical Image Analysis, 5(2), 143–156. 10.1016/S1361-8415(01)00036-6

Jenkinson, M., Bannister, P., Brady, M., & Smith, S. (2002). Improved optimization for the robust and accurate linear registration and motion correction of brain images. NeuroImage, 17(2), 825–841. 10.1006/nimg.2002.1132

Jung, E.-S., Dong, S.-Y., & Lee, S.-Y. (2019). Neural correlates of variations in human trust in human-like machines during non-reciprocal interactions. Scientific Reports, 9, 9975. 10.1038/s41598-019-46098-8

Krach, S., Hegel, F., Wrede, B., Sagerer, G., Binkofski, F., & Kircher, T. (2008). Can machines think? Interaction and perspective taking with robots investigated via fMRI. PLoS ONE, 3(7), e2597. 10.1371/journal.pone.0002597

Kriegeskorte, N., Simmons, W. K., Bellgowan, P. S. F., & Baker, C. I. (2009). Circular analysis in systems neuroscience: The dangers of double dipping. Nature Neuroscience, 12(5), 535–540. 10.1038/nn.2303

Krueger, F., & Meyer-Lindenberg, A. (2019). Toward a model of interpersonal trust drawn from neuroscience, psychology, and economics. Trends in Neurosciences, 42(2), 92–101. 10.1016/j.tins.2018.10.004

Lahnakoski, J. M., Glerean, E., Salmi, J., Jääskeläinen, I. P., Sams, M., Hari, R., & Nummenmaa, L. (2012). Naturalistic fMRI mapping reveals superior temporal sulcus as the hub for the distributed brain network for social perception. Frontiers in Human Neuroscience, 6, 233. 10.3389/fnhum.2012.00233

Lamichhane, B., Adhikari, B. M., & Dhamala, M. (2016). The activity in the anterior insulae is modulated by perceptual decision-making difficulty. Neuroscience, 327, 79–94. 10.1016/j.neuroscience.2016.04.016

Lee, J. D., & See, K. A. (2004). Trust in automation: Designing for appropriate reliance. Human Factors, 46(1), 50–80. 10.1518/hfes.46.1.50_30392

Madhavan, P., & Wiegmann, D. A. (2007). Similarities and differences between human-human and human-automation trust: An integrative review. Theoretical Issues in Ergonomics Science, 8(4), 277–301. 10.1080/14639220500337708

Microsoft. (2017). Dogs vs. cats dataset [Data set]. https://www.microsoft.com/en-us/download/details.aspx?id=54765

Moeller, S., Yacoub, E., Olman, C. A., Auerbach, E., Strupp, J., Harel, N., & Uğurbil, K. (2010). Multiband multislice GE-EPI at 7 tesla, with 16-fold acceleration using partial parallel imaging with application to high spatial and temporal whole-brain fMRI. Magnetic Resonance in Medicine, 63(5), 1144–1153. 10.1002/mrm.22361

Monfardini, E., Gazzola, V., Boussaoud, D., Brovelli, A., Keysers, C., & Wicker, B. (2013). Vicarious neural processing of outcomes during observational learning. PLoS ONE, 8(9), e73879. 10.1371/journal.pone.0073879

Mumford, J. A., Turner, B. O., Ashby, F. G., & Poldrack, R. A. (2012). Deconvolving BOLD activation in event-related designs for multivoxel pattern classification analyses. NeuroImage, 59(3), 2636–2643. 10.1016/j.neuroimage.2011.08.076

O’Doherty, J. P. (2004). Reward representations and reward-related learning in the human brain: Insights from neuroimaging. Current Opinion in Neurobiology, 14(6), 769–776. 10.1016/j.conb.2004.10.016

Parasuraman, R., & Manzey, D. H. (2010). Complacency and bias in human use of automation: An attentional integration. Human Factors, 52(3), 381–410. 10.1177/0018720810376055

Parasuraman, R., & Riley, V. (1997). Humans and automation: Use, misuse, disuse, abuse. Human Factors, 39(2), 230–253. 10.1518/001872097778543886

Pfeiffer, U. J., Schilbach, L., Timmermans, B., Kuzmanovic, B., Georgescu, A. L., Bente, G., & Vogeley, K. (2014). Why we interact: On the functional role of the striatum in the subjective experience of social interaction. NeuroImage, 101, 124–137. 10.1016/j.neuroimage.2014.06.061

Poldrack, R. A. (2007). Region of interest analysis for fMRI. Social Cognitive and Affective Neuroscience, 2(1), 67–70. 10.1093/scan/nsm006

Polich, J. (2007). Updating P300: An integrative theory of P3a and P3b. Clinical Neurophysiology, 118(10), 2128–2148. 10.1016/j.clinph.2007.04.019

Power, J. D., Mitra, A., Laumann, T. O., Snyder, A. Z., Schlaggar, B. L., & Petersen, S. E. (2014). Methods to detect, characterize, and remove motion artifact in resting state fMRI. NeuroImage, 84, 320–341. 10.1016/j.neuroimage.2013.08.048

Preuschoff, K., Quartz, S. R., & Bossaerts, P. (2008). Human insula activation reflects risk prediction errors as well as risk. Journal of Neuroscience, 28(11), 2745–2752. 10.1523/JNEUROSCI.4286-07.2008

Rauchbauer, B., Nazarian, B., Bourhis, M., Ochs, M., Prevot, L., & Chaminade, T. (2019). Brain activity during reciprocal social interaction investigated using conversational robots as control condition. Philosophical Transactions of the Royal Society B: Biological Sciences, 374(1771), 20180033. 10.1098/rstb.2018.0033

Ridderinkhof, K. R., Ullsperger, M., Crone, E. A., & Nieuwenhuis, S. (2004). The role of the medial frontal cortex in cognitive control. Science, 306(5695), 443–447. 10.1126/science.1100301

Rilling, J. K., & Sanfey, A. G. (2011). The neuroscience of social decision-making. Annual Review of Psychology, 62, 23–48. 10.1146/annurev.psych.121208.131647

Rilling, J. K., Gutman, D. A., Zeh, T. R., Pagnoni, G., Berns, G. S., & Kilts, C. D. (2002). A neural basis for social cooperation. Neuron, 35(2), 395–405. 10.1016/S0896-6273(02)00755-9

Rissman, J., Gazzaley, A., & D’Esposito, M. (2004). Measuring functional connectivity during distinct stages of a cognitive task. NeuroImage, 23(2), 752–763. 10.1016/j.neuroimage.2004.06.035

Rogers, D. A., Brooks, K. J., Finn, A., Schlesewsky, M., Ullsperger, M., & Bornkessel-Schlesewsky, I. (2026). The absent P3a. Performance monitoring ERPs differentiate trust in humans and autonomous systems. Psychophysiology, 63(7), e70354. 10.1111/psyp.70354

Rolls, E. T., Cheng, W., & Feng, J. (2020). The orbitofrontal cortex: Reward, emotion and depression. Brain Communications, 2(2), fcaa196. 10.1093/braincomms/fcaa196

Rosenberger, L. A., Eisenegger, C., Naef, M., Terburg, D., Fourie, J., Stein, D. J., & van Honk, J. (2019). The human basolateral amygdala is indispensable for social experiential learning. Current Biology, 29(20), 3532–3537.e3. 10.1016/j.cub.2019.08.078

Rosenthal-von der Pütten, A. M., Krämer, N. C., Maderwald, S., Brand, M., & Grabenhorst, F. (2019). Neural mechanisms for accepting and rejecting artificial social partners in the uncanny valley. Journal of Neuroscience, 39(33), 6555–6570. 10.1523/JNEUROSCI.2956-18.2019

Sanfey, A. G., Rilling, J. K., Aronson, J. A., Nystrom, L. E., & Cohen, J. D. (2003). The neural basis of economic decision-making in the ultimatum game. Science, 300(5626), 1755–1758. 10.1126/science.1082976

Shackman, A. J., Salomons, T. V., Slagter, H. A., Fox, A. S., Winter, J. J., & Davidson, R. J. (2011). The integration of negative affect, pain and cognitive control in the cingulate cortex. Nature Reviews Neuroscience, 12(3), 154–167. 10.1038/nrn2994

Uddin, L. Q. (2015). Salience processing and insular cortical function and dysfunction. Nature Reviews Neuroscience, 16(1), 55–61. 10.1038/nrn3857

Ullsperger, M., Danielmeier, C., & Jocham, G. (2014a). Neurophysiology of performance monitoring and adaptive behavior. Physiological Reviews, 94(1), 35–79. 10.1152/physrev.00041.2012

Ullsperger, M., Fischer, A. G., Nigbur, R., & Endrass, T. (2014b). Neural mechanisms and temporal dynamics of performance monitoring. Trends in Cognitive Sciences, 18(5), 259–267. 10.1016/j.tics.2014.02.009

Walbrin, J., Downing, P., & Koldewyn, K. (2018). Neural responses to visually observed social interactions. Neuropsychologia, 112, 31–39. 10.1016/j.neuropsychologia.2018.02.023

Wiese, E., Metta, G., & Wykowska, A. (2017). Robots as intentional agents: Using neuroscientific methods to make robots appear more social. Frontiers in Psychology, 8, 1663. 10.3389/fpsyg.2017.01663

Xu, J., Moeller, S., Auerbach, E. J., Strupp, J., Smith, S. M., Feinberg, D. A., Yacoub, E., & Uğurbil, K. (2013). Evaluation of slice accelerations using multiband echo planar imaging at 3T. NeuroImage, 83, 991–1001. 10.1016/j.neuroimage.2013.07.055

Yoshida, K., Saito, N., Iriki, A., & Isoda, M. (2012). Social error monitoring in macaque frontal cortex. Nature Neuroscience, 15(9), 1307–1312. 10.1038/nn.3180

